# Low-Frequency Tibial Neuromodulation Increases Voiding Activity - a Human Pilot Study and Computational Model

**DOI:** 10.1101/2024.05.17.594683

**Authors:** Aidan McConnell-Trevillion, Milad Jabbari, Wei Ju, Elliot Lister, Abbas Erfanian, Srinjoy Mitra, Kianoush Nazarpour

## Abstract

Despite widespread clinical adoption for disorders of incontinence such as overactive bladder, there remain unknowns surrounding the mechanism that underpins Tibial Nerve Stimulation (TNS). Current understanding suggests that TNS counteracts incontinence by the inhibition of brainstem and spinal cord activity. How this inhibition alters bladder function is not fully understood. We hypothesize that the supraspinal components of the system act as a high-pass filter, allowing voiding signals to proceed only when bladder filling reaches a critical level. Testing this hypothesis may explain how TNS is able to induce both an inhibitory and a little-explored excitatory effect on bladder activity in response to high-frequency (20 Hz) and low-frequency (1 Hz) stimulation, respectively. We performed a single-blinded trial in healthy human participants administered high and low-frequency Transcutaneous TNS. We also developed a computational model of the lower-urinary tract and control circuit to study the frequency-dependent effects of TNS. For the first time, we report a frequency-dependent effect of TNS via the ability to alter urge perception and up-regulate and down-regulate bladder activity, corroborating model predictions. These results provide a foundation for the development of targeted and effective TNS therapies, benefiting from *in silico* models. We hope that future clinical research will determine the efficacy of low-frequency TNS as a non-invasive treatment option for urinary retention.

**Significance Statement:** This work describes an experimental study supported by a novel computational model that for the first time captures the frequency-dependent effects of tibial neuromodulation on the urinary control system in humans. The findings of the work provide critically important evidence for the role of the brainstem as a filter, which may explain the little-explored excitatory effect of tibial neuromodulation on bladder activity. These results have considerable clinical implications in the treatment of urinary retention, a condition for which there are at present very few non-invasive treatment options available.

## Introduction

Micturition is regulated by a dynamic reflexive process that switches between periods of storage and release of urine held in the bladder. This cycle is mediated by a complex network of autonomic and somatic neurons in the central and peripheral nervous systems [1–3].

The tibial nerve has a significant influence on micturition [4, 5], because projections from the tibial nerve to the spine inhibit afferent sensory neurons [6–8] and potentially several midbrain nuclei, including the Periaquaductal Grey (PAG) and Pontine Micturition Centre (PMC) [9–11]. This inhibition has been shown to act through both classical and opioidergic mechanisms [12]. This differential mechanism of action likely explains the ability of the tibial nerve to inhibit both the nociceptive and non-nociceptive activity of the bladder [13].

The ability of the tibial nerve to down-regulate bladder activity has led to the formal adoption of tibial nerve stimulation (TNS) as a clinical intervention for the management of symptoms of the lower urinary tract [13, 14]. Non-invasive transcutaneous TNS (known as TTNS) is used as a clinically approved treatment for the management of incontinence and overactive bladder [15]. However, despite the widespread adoption of this technique in clinical practice, key aspects of the dose-related efficacy of TTNS remain unexplored. Specifically, preliminary feline evidence suggests that there may be a frequency-dependent effect of TTNS where low-frequency stimulation is able to up-regulate bladder activity, in contrast to high-frequency stimulation, which downregulates it [16–18].

We aimed to explore whether the preliminary evidence for a presently underexplored frequency dependence manifested as a functional effect in a human population and potentially shed light on a possible mechanistic explanation for the effect. We sought to achieve these goals with a two-pronged approach: conducting a pilot study in a healthy human population and developing a detailed computational model that can explain such frequency dependence *in silico*.

Therefore, we conducted a single-blind randomized control trial in a healthy adult population. In addition, we included a washout period to study if any observed effects linger in nature. With our computational model of the lower urinary tract and its controlling neural circuits, we simulated the bladder control network.

It is hypothesized that by varying only the frequency of applied TTNS, it will be possible to selectively up-or down-regulate bladder behavior in healthy adult humans. Moreover, we hypothesize that modeling the system computationally will allow elucidating the specific mechanism behind the effect.

We found that a frequency-dependent up-or down-regulation of bladder activity could be induced in a healthy human population. Furthermore, our computational findings corroborated these results and indicated that the effect may be mediated by brainstem-specific tibial projections and supported by spinal inhibition.

We propose a mechanistic framework for this effect based on the idea of filtering afferent sensory activity. These results reveal for the first time the presence of a frequency-dependent effect of TTNS in humans and provide a considerable foundation for future clinical research pertaining to the treatment of several disorders of the lower urinary tract.

## Methods

### Ethical Approval

The study was approved by the local ethics committee of The University of Edinburgh (ID: 977504). Forty-eight people were recruited under three experimental conditions. Before participation, they gave their full informed consent in writing.

### Study design

A graphical overview of the study can be seen in Fig. 1. In summary, the study was a single-blind design between subjects. Participants were pseudo-allocated to one of three groups such that the sample size across groups remained equally distributed (see below). Participants were not informed of this decision to ensure blinding was maintained.

**Figure 1.**
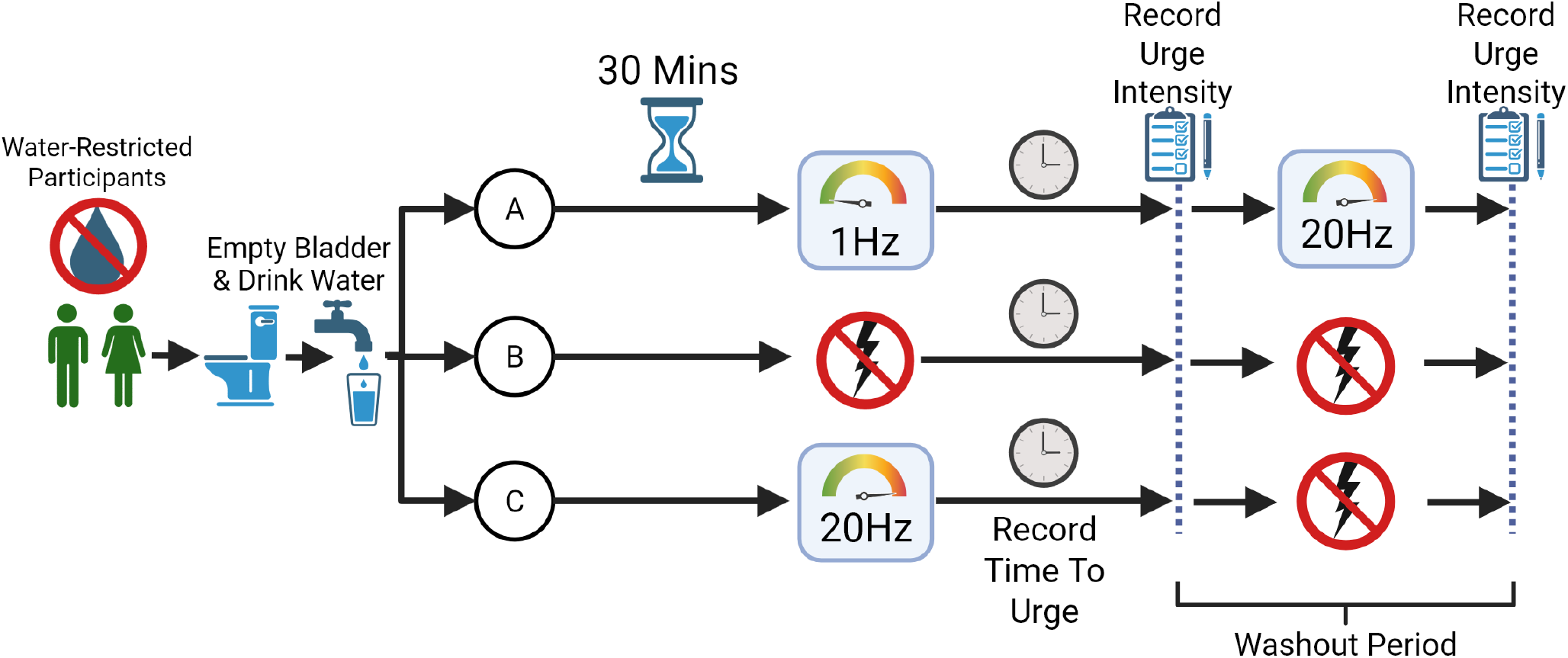
Experimental Study Methodology. Shown is a high-level overview of the study used to validate the frequency-dependent effects of TTNS predicted by the computational model. Participants were healthy adults (18+) asked to abstain from any nicotine/caffeine for 12 hours, and any fluids for 2 hours before the study. Upon arrival they were asked to ingest 750 ml water after emptying their bladder. A 30 min digestion period was employed before stimulation. Participants were pseudo-randomly allocated between groups, and were blind to the condition.

To ensure a relatively stable bladder baseline across the sample, several measures were taken. First, participants were asked to restrict their fluid intake for two hours and caffeine or nicotine for 12 hours before the beginning of the experiment - caffeine or nicotine affect urination [19, 20]. Second, participants were asked to empty their bladder as much as they were reasonably able upon arrival, before ingesting 750 ml of water.

Participants underwent a 30 minute “digestion period” during which they were able to undertake any activities they wanted as long as they remained seated. During this period, they were prepped for neurostimulation: Any leg hair was shaved from the ankle and lower leg using a sterile disposable razor, and the stimulation site was disinfected using a 70% isopropyl alcohol solution to remove contaminants.

To ensure that the objective outcome of the study could be recorded in all cases, after the digestion period, participants were told to inform researchers when they first felt the urge to urinate. Electrodes (*Med-Fit 50mmx50mm Hydrogel Electrodes*) were applied to the right leg 1 *cm* posterior to and 10 *cm* superior to the medial malleolus in accordance with established TTNS protocols [13]. Participants received stimulation (*Digitimer DS7A High Voltage Constant Current Stimulator*) until they felt the urge to urinate. The specific parameters of the stimulation differed according to group allocation (see fig. 1):

- Group A: 1 Hz, 200 *µs* pulse width, motor threshold.
- Group B: Control group, without stimulation, though they still underwent skin preparation.
- Group C: 20 Hz, 200 *µs* pulse width, motor threshold.

In all cases, any applied stimulation was monophasic and administered at a standard voltage (max 300V, see supplementary materials) with a current sufficient to induce flexion or fanning of the big-toe. The current was slowly increased until the first sign of this motor activity was observed.

Upon reporting the urge to urinate, the time-elapsed was recorded and the participants were asked to rate how intense they felt their urge according to a validated questionnaire [21] (see Supplementary Materials).

### Washout Period

To analyze the presence of lingering effects and to explore whether the hypothesized excitatory effect was reversible, an additional washout period was included in the study methodology, as shown in Fig. 1). After reporting the urge to urinate, participants were asked if they would be willing to take part in an additional optional experiment. If they agreed, participants underwent an optional 10 minute extension to the study where:

- Group A received 20 Hz stimulation, 200 *µs* pulse width, motor threshold.
- Groups B & C were asked to remain seated for 10 minutes and no stimulation was applied (the electrodes were disconnected from the stimulator).

To maintain the blinding of participants to their initial group allocation, participants in groups B and C were told that the optional experiment would simply involve sitting without the electrodes on for ten minutes. At this point the electrodes were removed from the ankle (to ensure they were aware no stimulation was applied) and the washout period initiated for these groups. Individuals in group A were informed that the optional experiment would involve administering a different kind of stimulation which would feel different.

After this washout period, a final urge intensity score was obtained before the end of the study.

### Model Topology

The simulated network was composed of a set of interconnected neuronal units each comprising 100 individual neurons modeled in terms of conductance, or Poisson point processes that generated simulated action potentials (see Neuronal Model). The topology of the model was adapted from previous work that simulated reflex voiding [22]. The circuit remained broadly unmodified, with the exception of several key modifications, including simulation of the tibial nerve and its projections (see Fig. 2). We focus on normal bladder function; therefore, the nociceptive bladder afferents were not included in our model. To simulate the effects of opioidergic and classical inhibition, neurons and synapses were represented in terms of ionic conductance. The biophysical model used to represent the bladder was adapted from [23](see Bladder Model).

**Figure 2.**
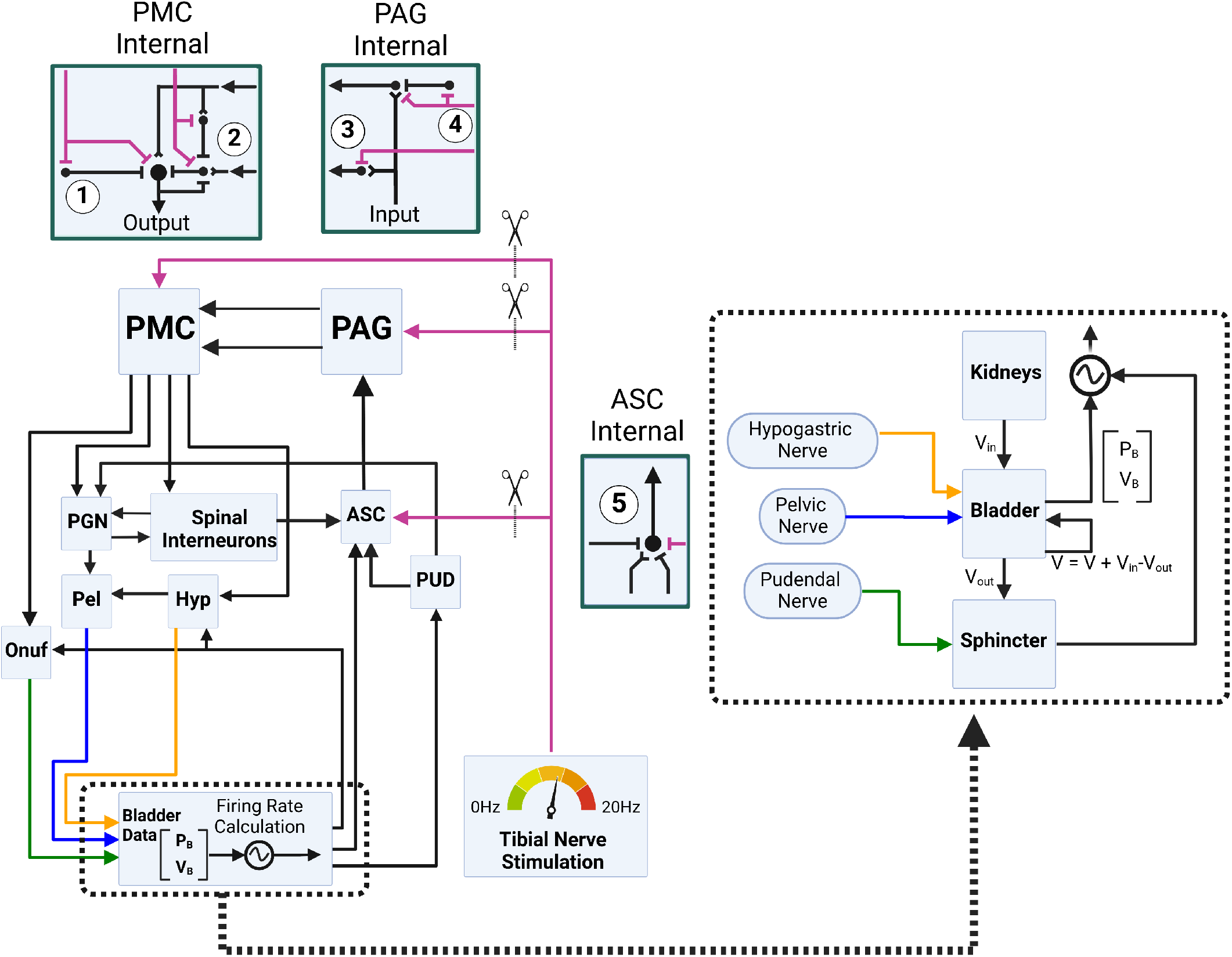
Overview of the Computational Bladder Control Model. **A**. Shown is a block diagram of the simulated neuronal circuit and bladder model. The model used a modified biophysical representation of the bladder produced previously (dashed box, [23]) which used the firing rates of the pelvic (blue), hypogastric (orange), pudendal (green) efferents to calculate bladder state. Tibial input used to modulate the circuit shown in purple. PMC: Pontine Micturition Centre, PAG: Periaquaductal Grey, PGN: Preganglionic Bladder Neurons, ASC: Ascending Interneurons, PUD: Pudendal Afferent, Onuf: Onuf’s Nucleus, Pel: Pelvic Efferent, Hyp: Hypogastric Efferent. Scissors represent possible severed projections. Labels 1 - 5 represent key regions of tibial modulation using opioidergic (1 - 4) or classical (5) inhibitory mechanisms.

The tibial nerve was modeled to project to second-order sensory afferents (via classical inhibitory synapses) and all neuronal units within the PAG and PMC (via opioidergic inhibitory synapses) as shown in Fig. 2. To provide fine control over the level of tibial modulation applied to the system, the firing rate of the neuron and its projections were clamped and, if required, could be severed by disconnecting synaptic junctions. This ensured that while in a severed state, the neurons though still modeled had no impact on post-synaptic targets.

### Bladder Model

The model used a biophysical representation of the bladder adapted from the work published by Lister et al. [23] modified to allow integration with the conductance-based representation of the micturition control circuit used in the present work. In summary, for each step of the simulation (*t*), the firing rates of three key efferents: Pelvic, Hypogastric, and Pudendal were used to calculate the internal activation constants - *ω*_*e/i/s*_ - which governed detrusor excitation/inhibition, and sphincter activation respectively. From these values, the bladder state could be calculated and the internal detrusor pressure (*P*_*B*_(*t*)) used to update the afferent neuronal activity (the main input for the control circuit; see Fig. 2). To ensure computational efficiency, the sampling rate of the model was set at 50 Hz, with each step of the simulation (Δ*t*) representing a 20-ms increment. A detailed description of the model can be found in [23].

### Computational Model Fitting

To fit the synaptic weights of the model (see Supplementary Materials), we utilized real bladder pressure and neural data from a previously published work (see Jabbari et al. 2019[24]. In brief, the training data was obtained from intact adult male Wistar rats. The dataset was comprised of bladder pressure and volume information, and extracellular neuronal recordings obtained from between the L6 to S1 spinal cord.

### Neuronal Model

The model was built using the Brian2 package for Python [25]. The relationship between the pelvic afferent firing rate and bladder pressure was calculated according to an empirically derived relationship determined previously [26] where the firing rate of the primary bladder afferent at each moment (*F*(*P*_*B*_(*t*))) was defined as:

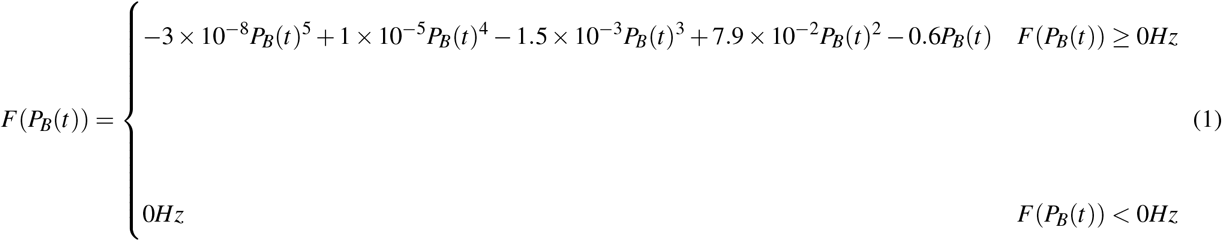

All neuronal units, with the exception of primary bladder afferents and the tibial nerve, were simulated using a Conductance-Based Adaptive Exponential Integrate-and-Fire (CAdEx) model [27] modified to include opioidergic inhibition. Through the inclusion of an additional input current (*I*_*ap*_) neurons could be made tonically active where required. This was particularly important in the simulation of PAG and PMC, where inhibitory feedback mechanisms reliant on tonic activity were present.

The tibial nerve and primary bladder afferents were simulated as Poisson point processes which generated simulated neuronal spikes at a pre-specified rate. Doing so allowed the firing rate of the pelvic afferent to be set according to the aforementioned mathematical relation and the firing rate of the tibial nerve to be clamped to a specific value (representing external stimulation of the nerve).

Each *unit* within the network was composed of 100 simulated neurons connected via random synapses (where no autapses were permitted) to introduce noise into the system.

Membrane potential (*v*) was defined as

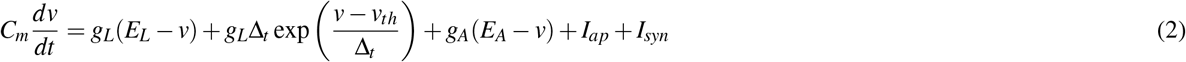

where *C*_*m*_ was the membrane capacitance; *g*_*L*_ the leakage current & *E*_*L*_ its associated reversal potential; *g*_*A*_ was the adaption current and *E*_*A*_ its reversal potential; *I*_*ap*_ the input to tonically firing neurons; and *I*_*syn*_ the input from synaptic connections. Also included were terms that govern the action potential (*v*_*th*_ and Δ_*t*_). As with the original definition proposed buy Gorski et al,[27], *g*_*A*_ was defined as:

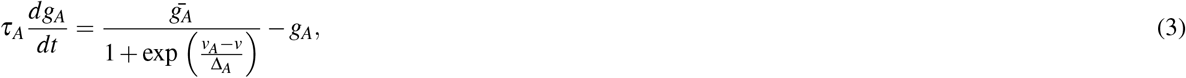

where the rate of change of the current was determined by a rate constant (*τ*_*A*_), the subthreshold adaption parameters (*v*_*A*_ and Δ_*A*_), and the maximum subthreshold adaption conductance 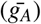. The model also included a post-spike reset mechanism, which too remained unchanged.

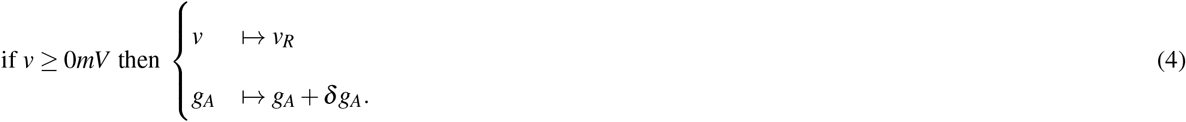

Upon spiking, the membrane potential was reset to its resting state (*v*_*R*_), and the adaption current increased (*δg*_*A*_). The purpose of the adaption current was to allow modeling of the spike-frequency adaption. Tonically active neurons did not display spike-frequency adaption to ensure that their activity remained consistent.

### Tibial Nerve Stimulation Analysis

To computationally analyze the impact of tibial modulation, simulations were performed in a range of modulatory states. Tibial modulation was applied at 21 different frequencies (0-20 Hz, in 1 Hz increments, *N*_*repeats*_ = 10 in each condition) to a simulation of 500-second duration (where each 500 second simulation was run separately). Any aspect of simulated bladder function or circuit activity could be recorded during these simulations. Where neuronal firing rate activity was analyzed, instantaneous firing rates for each recorded time step were smoothed (1st-order Butterworth filter, with a 1 Hz cutoff). To analyze the effect of specific tibial projections, projections to regions of interest could be severed as necessary to prevent the formation of synaptic connections.

## Results

### Participants Display a Frequency-Dependent Sensory Response to TTNS

Before statistical analysis was conducted, several datapoints had to be removed from the dataset due to:

- A failure to abstain from caffeine/nicotine before participation (*n* = 1, Group A)
- Presence of an ongoing treatment for relevant co-morbidities (*n* = 1, Group A)
- Several participants reporting the urge to urinate before the end of the digestion period, preventing administration of neuromodulation (*n* = 3, *n*_*GroupA*_ = 1, *n*_*Groupb*_ = 2)

Unfortunately, exclusion of these participants did lower the statistical power of the analysis, resulting in a final final sample distribution of *N*_*A*_ = 14, *N*_*B*_ = 14, *N*_*C*_ = 15. The non-normality and relatively small sample size of this distribution required that non-frequentist statistical techniques be used.

Despite this, the results of the study were nonetheless promising. Analyzing the effects of the intervention, it was clear that there was a differential effect of stimulation frequency on urge-onset time. To account for the small sample size, and non-normality of the dataset, a Bayesian Robust Linear Regression was conducted, rather than a typical frequentist approach. Given the potential for outliers arising as a result of the sample size, a Student’s t distribution was selected during likelihood calculation (*N*_*draws*_ = 12, 000). The model indicated that group A (low-frequency TTNS) had a mean time to first urge of 937 seconds (*HDI*_3%_ = 575.88*s, HDI*_97%_ = 1291.88*s*, see fig. 3A). Posterior distribution comparison indicated an 89.28% probability that individuals exposed to low-frequency TTNS experienced urge-onset more rapidly than placebo control (group B, *mean* = 1277.7*s, HDI*_3%_ = 880.68*s, HDI*_97%_ = 1648.17*s*). As detailed in fig. 3, also indicated that group C (high-frequency TTNS) displayed a delayed urge onset compared to placebo control (*mean* = 2301.93*s, HDI*_3%_ = 1931.67, *HDI*_97%_ = 2642.83*s*) with posterior analysis indicating a 99.94% chance that high-frequency intervention delayed urge onset in individuals within group C, when compared to those in group B. All parameters indicated excellent model convergence 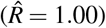. These results indicate the likely presence of a promising frequency-dependent trend, however additional analysis is critical as the sample size remains too small to make any certain claims at this stage.

**Figure 3.**
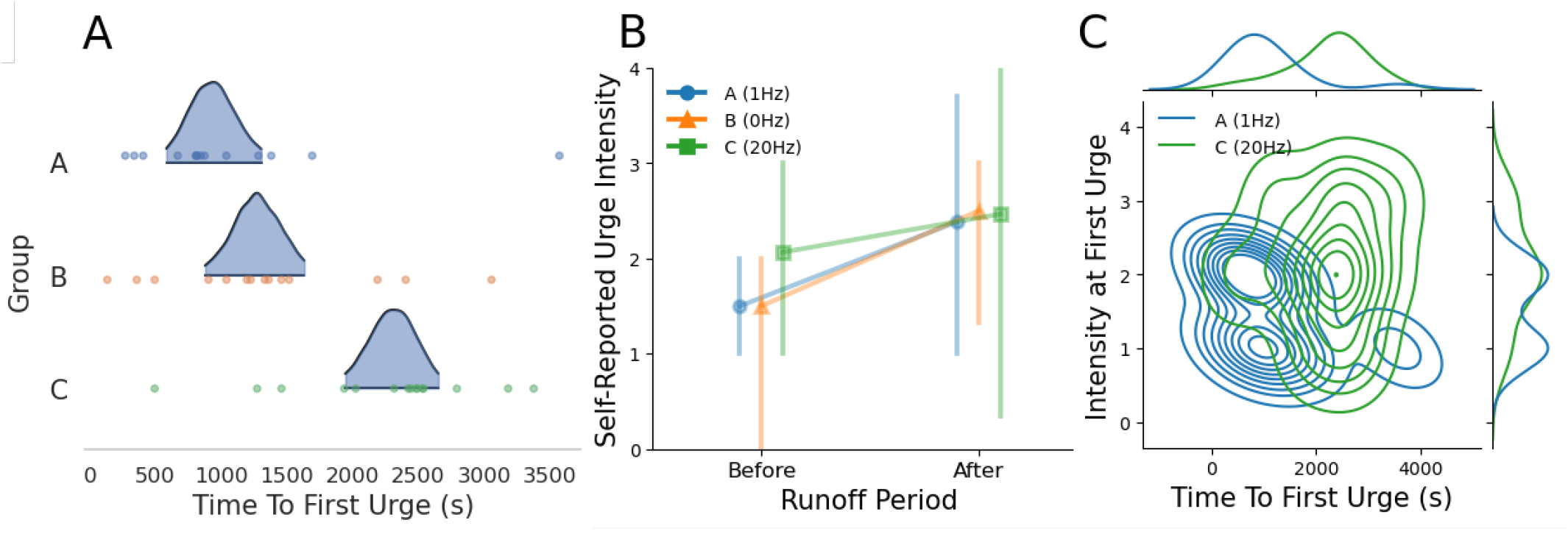
Experimental Study Results. **A**: effects of low (1 Hz, Group A), high (20 Hz, Group C), and placebo (0Hz, Group B) TTNS stimulation on the time elapsed to first sensation of urge. Shown are raw-data points and Bayesian posteriors for each group.**B**: effects of the washout period on self-reported urge intensity. Error-bars = *±*95% Confidence Interval. Urge intensity self-reported on a scale of 0–4 (see methods), washout period was 10 minutes in duration. **C**: bivariate and univariate kernel-density estimate plot displaying the relationship between the time elapsed before participants reported the urge to urinate (i.e., the unit reported in panel A), against the self-reported intensity of this urge at (i.e., how strongly participants felt this initial urge, the data reported in panel B before the additional 10-minute washout period).

Nevertheless, an additional Region of Practical Equivalence (ROPE) analysis was conducted to assess whether the difference in urge-onset time between groups A and B (which would indicate a novel effect of TTNS) were clinically relevant. A range of *±*60*s* was selected as the region of practical equivalence for the analysis. Doing so revealed that 7.49% of the posterior distribution of the difference in urge onset fell within the region of equivalence indicating a considerable likelihood of potential clinical relevance (see Supplementary Materials)

### TTNS Has Little Frequency-Dependent Effect on Perceived Urge Intensity

Of all experimental participants (including those removed from statistical analysis) only one individual declined the offer to take part in the washout period, resulting in a participation rate of 98%.

A factorial ANOVA suggested there was a significant main effect of the washout period (*F*_(1,79)_ = 17.625, *p <* 0.00001) but not group allocation (*F*_(2,79)_ = 1.276, *p* = 0.28) on urge intensity. In addition, there was no significant interaction between the factors reported (*F*_(2,79)_ = 1.076, *p* = 0.35). In all cases, there was an increase in the average urge intensity reported by the participants before and after the washout period (see Fig. 3B). At the end of the washout period, the three conditions finished with a similar intensity score. However, the variability within this data makes any robust inferences impossible.

There was a small but obsertable relationship between urge intensity and sensory response (i.e., time elapsed to urge sensation). While urge intensity at the first reporting point (i.e., before the washout period) was similar across groups, those in the excitatory condition reported the sensation earlier compared to those in the inhibitory condition (see Fig. 3C).

### Normal Bladder Activity May be Computationally Modeled

Under baseline (unmodulated) conditions, the behavior of the model was promising. The model accurately simulated normal bladder function and the associated efferent nervous activity. Specifically, it was able to reproduce the expected filling and voiding cycle (see Fig. 4 top trace). During the filling phase, the efferent projections associated with urine storage (hypogastric and pudendal) were tonically active, while the pelvic projections associated with voiding were silent.

**Figure 4.**
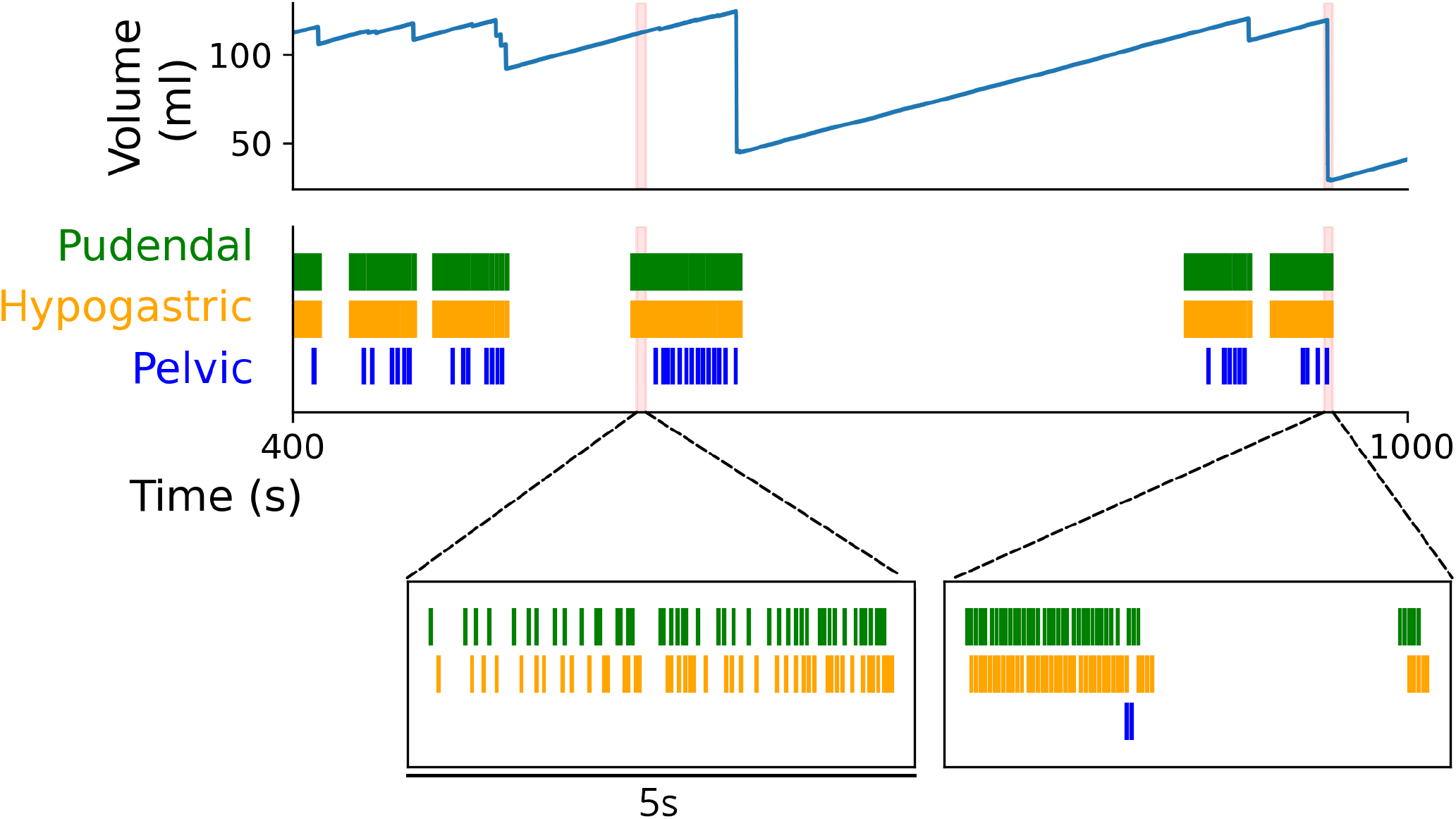
Behavior of the Simulated Bladder Model. Shown is the bladder behavior (top trace) and associated efferent neuronal activity (middle trace) recorded from a 1000-second duration simulation. Blowout boxes contain 5 second windows of the full recorded activity (highlighted in red) during filling (left box) and voiding (right box). Each raster trace was obtained from a randomly selected neuron within the Pudendal, Hypogastric, and Pelvic units (*N*_*neurons*_ = 100 in each case).

In contrast, during void events, bursts of pelvic efferent activity silenced the hypogastric and pudendal projections. This change in activity was associated with a decrease in bladder volume, indicative of a switch in the system from a storage state to voiding (see fig. 4). These results suggest that the model was able to mimic the complex switching behaviors required to mediate the behavior of the lower urinary tract, allowing it to serve as an accurate baseline upon which to test the impact of modulation.

Though the model does display the ability to accurately capture bladder-like behaviour it should be noted that there were several limitations that may limit its generalisability. The model displayed a much smaller bladder capacity (mean 112*ml, ±*11.6) than observed physiological values in humans (300 − 500*ml*[28]). Additionally, the model displayed poor voiding efficiency. Under normal (unmodulated) conditions the average voiding efficiency was 14.5% (*±*31.4%). The low average efficiency coupled with the relatively large variance limits the direct physiological comparisons that may be drawn from the model.

These limitations are likely due to the training data used during model fitting. The original dataset was obtained from male Wistar rats[24]. The average capacity across the dataset was 0.7*ml* (*±*0.09). Moreover, it also displayed a similarly poor voiding efficiency throughout (mean=9.54%, *±*2.59) - likely due to the tightly constrained nature of the recorded bladder behaviour. While the reduced capacity and voiding efficiency of the training dataset were partially overcome (as the performance of our model outperformed in these metrics slightly) the limited generalisability of the model should be kept in mind when interpreting the results.

Despite this limited generalisability the model never nevertheless serves as a more than acceptable foundation for drawing proof-of-concept conclusions and examining potential mechanistic explanations for tibial nerve simulation.

### Computational Modeling Confirms the Frequency-Dependent Effects of Tibial Nerve Stimulation

Analyzing the effects of TNS on the computational model revealed an intriguing trend. Under baseline conditions, the model produced a cycle of storage and voiding in accordance with expectations from previously obtained animal data (see [24], Fig. 5A). Applying high-frequency (20 Hz) stimulation to the system completely inhibited the voiding action (according to the expected effect of TNS on bladder behavior). However, if low-frequency (1 Hz) TNS was applied to the system, voiding occurred earlier (baseline mean=426*s, ±*14.6; low-frequency mean=410*s, ±*9.9) and with greater voiding efficiency (baseline mean=9.54%, *±*2.59; low-frequency mean=41.44%, *±*21.72) causing larger drops in stored volume for a given voiding event (see fig. 5A and supplementary materials).

**Figure 5.**
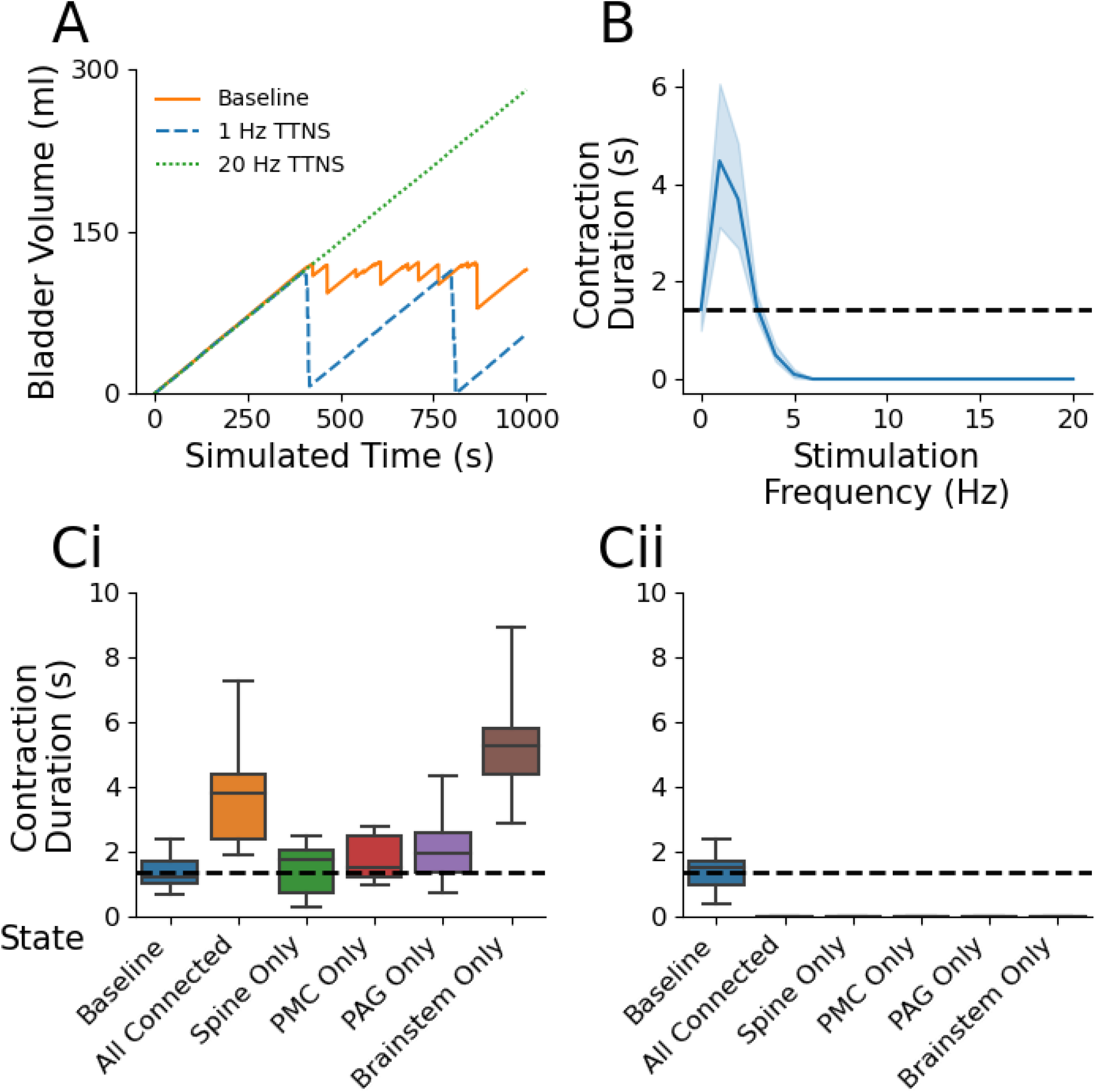
Computationally Modeling TTNS. **A**. Effects of low (1 Hz) and high (20 Hz) frequency TTNS on simulated bladder function compared to unmodulated (control) conditions. **B**. Frequency dependent effects of TTNS on bladder contraction. Shown is the average total bladder contraction duration for a 500s simulated period under 21 different stimulation frequencies (0-20 Hz, in 1 Hz increments, *N*_*repeats*_ = 10). Errorbar= *±*95%*CI*. A contraction duration of zero ms indicates that voiding was completely inhibited (i.e., no voiding events occurred during the simulated period of time) **C**. Effects of disconnecting specific tibial-nerve projections on total contraction duration (for all voids over a 500s simulation period) under low-frequency (1 Hz, **i**) and high-frequency (20 Hz, **ii**) conditions. In both cases, baseline behavior represents the unmodulated behavior of the model (i.e., where all tibial-nerve projections remained intact *and no tibial-stimulation was applied*). In contrast, the “all connected” conditions describe a model configuration where all tibial-nerve projections remained intact *and with tibial-stimulation applied at 1 Hz (i) or 20 Hz (ii). N*_*repeats*_ = 10 in each condition.

To do so, the research team conducted a further exploration of the system during low-frequency TNS. To guide this exploration an initial hypothesis was generated. In theory, to drive any increase in voiding activity *without* considerably changing the time elapsed before voiding (i.e., the time of void-onset) one or more of the following changes may be occurring:

- The intensity of bladder contraction is increased
- The duration of bladder contraction is increased
- The system-level mechanisms that control urine storage are being inhibited.

Further analyzing the behavior of the system under a range of different stimulation conditions, the research team discovered a frequency-dependent relationship between TNS and contraction *duration*. For a given simulation, at low frequencies the average effective duration of contraction – i.e., bladder contractions that successfully induced a voiding event were increased significantly, while at high frequencies they were eliminated completely, resulting in a contraction duration of 0ms (as voiding became impossible, see fig. 5B). These findings confirm the presence of a frequency-dependent effect of TNS on bladder behavior and indicate that the low-frequency effect may be driven by the extension of bladder contraction, causing a greater intensity of voiding for a given simulation run.

### Brainstem Targeting Projections Mediate Low-Frequency Excitatory Effects

As the duration of contraction was highlighted as a measure of interest by the initial computational analysis, we sought to understand the nature of this frequency-dependent effect by analyzing the specific projections that may mediate it. To do so, an analysis of bladder contraction was performed in a range of connective states (where specific tibial nerve projections were severed) during low-and high-frequency stimulation.

A one-way ANOVA revealed a significant effect of the connective state on the total duration of contractions (i.e., across all voiding events) over a 500s simulation during low-frequency (*F*_(5,54)_ = 16.975, *p <* 0.0001) and high-frequency conditions (*F*_(5,54)_ = 55.269, *p <* 0.0001).

From these results, it is clear that brainstem, rather than spinal cord, targeting projections are necessary for the manifestation of the low-frequency effect, though it cannot be said at this stage how each module contributes to the effect (as both were required to produce a large-magnitude increase in contraction duration). The effect size of the PAG only condition was larger than that of the PMC condition providing some evidence that this region may be playing a greater role in the process however further research is required to disentangle the individual effects of each region. Intriguingly it appears that the inclusion of spinal-targeting tibial projections has a pronounced influence on voiding efficiency, with the elimination of these projections (leaving only brainstem projections intact) causing a more consistent increase in voiding efficiency.

### The Mechanism Underlying High-Frequency Inhibition Remains Unclear

In contrast, the combination of projections that underlie high frequency stimulation is more opaque. Bladder contractions were extinguished by any combination of spinal or brainstem targeting tibial projections when 20 Hz TNS was applied. (see Fig. 5Cii). This finding makes it difficult to pinpoint the exact mechanism that underlies high-frequency inhibition of bladder activity (as all tested configurations were able to induce the effect). These findings require further analysis, but nevertheless provide additional evidence for a clear inhibitory effect of high-frequency TNS on bladder function.

## Discussion & Conclusion

### Experimental Study Findings

The results of the present study demonstrate that TTNS is able to impart a frequency-dependent effect on bladder function. They corroborate preliminary animal data within the literature [16–18] and for the first time demonstrate evidence for a frequency dependence in humans. The results suggest that the main effect of the intervention on the healthy population was sensation. Changes in the time taken for participants to experience the urge to urinate were the primary change in bladder behaviour, in contrast to the effects on urge intensity which were minimal, though significant variance within the data makes any conclusions difficult. Although 20 Hz stimulation appeared to increase the average intensity of the urge after treatment, this is unlikely to indicate the presence of any lingering effect. Rather, it is more likely that this difference compared to 1 Hz stimulation was due to more time elapsing before participants reported any feeling of urgency (see Fig. 3C). In other words, while a similar intensity was reported upon feeling the urge to urinate, the time taken to produce this feeling could be modulated - either reducing it (via 1 Hz stimulation), or increasing it (via 20 Hz stimulation). This result is surprising, as it would be expected than any modulation in one aspect of afferent bladder sensation (the urge onset) would reasonably drive a change in the other (the urge intensity). While this juxtaposition undoubtedly warrants further exploration it is possible to hypothesise a potential explanation of the effect. As will be shown by the results of the computational modeling it may be the case that any afferent sensory changes observed are secondary to the primary effect of TTNS - the modulation efferent bladder activity. TTNS has been shown to alter the activity of a number of regions of the brainstem and cortex associated with bladder function[29] in different ways dependent on bladder volume[30] suggesting a complex state-dependent sensory effect. Though not explored in the present publication it is possible these secondary effects (i.e., those outwith the PAG and PMC targets modelled here) influence the conscious perception of bladder sensation in a complex and temporally specific manner.

### Computational Findings

The limitations of the pilot study are somewhat remedied by the results of the computational modeling work which provide considerable mechanistic context to the experimental study. The unmodulated behavior of the model was promising, suggesting that it was able to accurately mimic the complex switching behavior employed by the micturition control system[2]. The system produced tonic storage-related efferent activity indicative of a guarding reflex during periods of lower bladder volume which transitioned to void-promoting pelvic activation and silence of storage-related activity at appropriate times. The switch in the circuit state aligned with voiding events in the data. These results suggest the model is able to capture both aspects of micturition (upper circuit activity and lower-bladder behavior) providing an accurate overview of the system for further research. Although the average bladder capacity of the model was lower than would be expected *in vivo* (simulated capacity was approximately 100-150ml compared to a physiological range of 300-400ml [31]) the accuracy of the high-level behavior of the system allowed the effects of TTNS to be studied *in silico*. The impact of TTNS on the system was clear: By varying solely the frequency of stimulation, it was possible to up-or down-regulate simulated bladder activity. This was in line with the hypothesis stated in the article, confirming the presence of a frequency-dependent effect highlighted by the experimental study. Though these results are promising there are several key limitations to our computational approach that should be kept in mind when interpreting any results. As previously mentioned the model displayed a bladder capacity and voiding efficiency considerably lower than normal human physiological ranges. Though this may be explained by the training data used during the fitting process it nevertheless reduces the physiological generalizations that may be drawn from the model. Moreover, the model did not simulate nociceptive afferent projections. Therefore, at present it cannot be used to test the efficacy of TTNS on a pathologicaly active lower urinary tract system. As such, care should be taken when directly comparing the computational findings to biological reality as they cannot directly inform us of any therapeutic clinical effects. Rather the present findings should be viewed through the lens of a systems level proof-of-concept approach. Nevertheless, even with a relatively narrow system-level perspective, our results still provide further evidence and a potential mechanistic explanation for an under-explored excitatory effect. These findings are more than sufficient to serve as an initial foundation for future clinical work.

### Joining the Two

Though these results are evidently promising, when the computational and human findings are taken together, they appear to paint different pictures of the frequency dependent effect. The pilot study indicates a primary effect of the intervention on afferent sensation (with urge sensation being the primary measure) while the computational model primarily indicates an efferent effect through a change in contraction duration (though there was a minor afferent effect present). Analysing the computational results, it is tempting to assume the primary effect of TTNS was on efferent behaviour as the change in contraction duration observed *in silico* was considerably greater than the equivalent shift in void onset time, mirroring the relatively small effect size seen in the pilot study where 1 Hz stimulation was found to induce the urge slightly earlier, but not at a considerably greater sensory magnitude. However, the relatively small sample size of the pilot study, and the simplified afferent model (which did not include nociceptive C-Fibres) preclude this conclusion at the present time.

The two measures are not unrelated. Both pertain to different aspects of bladder function, and as such further research must critically asses how the two outcome measures are connected, as the present work indicates that both may be affected by the TTNS in a frequency-dependent manner (though to varying degrees of functional efficacy). Nevertheless, these findings undeniably indicate to the presence of some form of complex frequency-dependent effect of TTNS on bladder related function

### Clinical Relevance and Research Direction

The future clinical research directions of this frequency dependence are particularly important when considering the treatment of non-obstructive urinary retention (NOUR). Currently, the treatment of NOUR remains rather limited. Invasive sacral stimulation is the only approved neuromodulatory treatment option available to patients [32]. Previous attempts to use TTNS to treat NOUR have returned mixed results [33, 34]. However, this may be due to the use of inappropriate stimulation parameters. Previous research did not alter the stimulation parameters of the standard interventions used in the treatment of incontinence and, as such, may have in fact had a deleterious effect on NOUR symptomatology.

Moreover, at present, TTNS is typically applied to treat incontinence prophylactically, with weekly sessions promoting long-term improvement in symptomatology [35, 36]. However, these findings suggest that if TTNS is to be applied in the treatment of urinary retention, it may be more effective to administer the treatment in an acute manner, i.e., at the point of voiding. Doing so could up-regulate the bladder and facilitate unassisted voiding. If shown to be effective when implemented in this way, it could serve as an effective means of reducing the dependency of urinary retention patients on catheters, reducing the rates of urinary tract infections and improving overall quality of life [37, 38].

In addition to providing evidence for therapeutic value in the treatment of retention, the present work may also shed light on the mechanistic underpinnings of this frequency-dependent effect. Our results indicate that low-frequency excitation is mediated by brain stem-specific projections. However, this does not preclude an influence of spinal projections. We found a minor but observable influence of spinal cord projections on voiding efficiency. Keeping spinal projections intact during low-frequency stimulation, seemingly reduced the magnitude of the observed increases in voiding efficiency (when compared to the same experiment run with only brainstem projections in place). It cannot be said at this time what the specific interaction between spinal and brainstem projections but it is clear there is a complex relationship between the two which should be explored in future work.

Regardless of the interaction between the spinal and brainstem projections, it may be possible to characterise the clear role of the brainstem in low-frequency excitation from a system-level perspective by characterising the region as a filter. The inhibitory feedback mechanisms within the PAG and PMC (see Fig. 2 labels 1–4) allow the modules to act as a set of high-pass filters connected in series, preventing efferent void signaling until a sufficient afferent firing rate is reached. Low-frequency TNS can induce an excitatory effect by inhibiting feedback mechanisms within the PAG, reducing the threshold of the filters, thus allowing a smaller afferent signal to pass through to the PMC (the primary micturion control region [2]) triggering void-promoting efferent activity.

While, inhibition of the PMC or PAG alone did not result in the same magnitude of excitation, this filtering hypothesis may explain the slight difference in both contraction duration and voiding efficiency between the two conditions. In both cases, contraction duration and voiding efficiency were greater when the PAG was targeted over the PMC. As the first filter in series (the PAG) it is possible that when left fully operational (i.e., where only intact PMC targeting tibial projections were active) it is capable of eliminating afferent activity before it can reach the PMC. In this configuration, the inhibitory state of the PMC matters little.

Although this filtering hypothesis effectively explains the relationship between spinal and brainstem projections during low-frequency TTNS, it cannot explain the complex interactions that may be at play during high-frequency TTNS. The results from the present study were unclear in this respect, as all conditions resulted in the same extinguishing of bladder behaviour making the formation of any explanation difficult. It is unclear if high-frequency inhibition occurs at the spinal level, silencing afferent activity before it reaches the filters (Fig. 2, label 5), or at the level of the brainstem, silencing communication between the modules that allows the production of void-promoting efferent activity (Fig. 2, labels 2–4) or even the efferent activity itself (Fig. 2, label 1). Preliminary evidence from the literature paints a picture of a complex relationship between spinal and brainstem tibial projections. Specifically, TTNS has been shown to have a directly inhibitory effect on spinal activity [8] that is nevertheless insufficient to induce bladder changes in cases of spinal cord injury [4]. Although this effect may be due to a rewiring of the bladder control system (as there is evidence that early application of TTNS has some effect [39]), coupled with the results from the present work it is clear there is a need for further research to explore the spinal and supraspinal aspects of the system interact to inhibit bladder function during high-frequency TTNS.

Future research should primarily focus on confirming if these system-level proof-of-concept results generalise from an *in silico* demonstration to real-world effects in a pathological population. Specifically, subsequent work should maintain a clear focus on exploring the clinical efficacy of low-frequency TTNS as a treatment option for urinary retention. Specifically, it would be prudent to determine if administration of the intervention in an acute manner, at a lower frequency, induces an objective, measurable change in bladder physiology. If followed through to completion, this research direction may remedy the mixed results seen previously. Additionally, these findings may contribute to the ongoing development of a wearable neuromodulatory treatment device which may provide a user-friendly method of administering the intervention [40].

## Conclusion

Despite some limitations surrounding the methodology of the experimental study and the need for further research in this direction, the present findings nonetheless provide vital evidence for a frequency-dependent effect of TTNS in human beings. Moreover, they suggest a complex and not-necessarily solely brainstem-specific role of the tibial nerve in bringing about this unexpected, but critically important excitatory effect. We propose that these findings would lay the groundwork for a number of future research efforts which may elucidate the precise neural mechanisms underlying the complex effects of TNS on bladder function in both healthy and diseased states.

## Code availability

The custom code used for the analysis in this study is available at[41]. The authors request that any individuals using the code as part of further research cite the present work.

## Author Contributions

A.M.T and K.N. conceived the study, A.M.T. constructed the computational model, and analyzed the results; M.J. and A.E. collected and processed the bladder data used to fit the computational model and run simulations in the present study; W.J. assisted with the recruitment and running of the study; E.L. created and provided the biophysical bladder simulation used in the computational model; A.M.T and M.J. produced the figures used in the publication; K.N. and S.M. provided research guidance and advice; All authors reviewed the manuscript.

## Acknowledgments

This project is funded in part by the University of Edinburgh and the Engineering and Physical Sciences Research Council (EPSRC) grant number EP/W031493/1.

## Supplementary Materials

### ROPE Analysis

As part of the pilot study statistical analysis, a Region of Practical Equivalence test was conducted. A region of practical equivalence of *±*60*s* was selected as an appropriate interval. Detailed in Supplementary fig. 4 is the result of the analysis, showcasing a 7.49% overlap in the distribution of differences when comparing groups A, and B.

### Bladder Urgency Survey

As part of the human study, participants were asked to rate how intense they felt the urge to urinate. This rating was adapted from a validated questionnaire typically used to rank urgency perception as a measure of urological dysfunction [21] (see Supplementary Fig. 1).

### TTNS Stimulation Parameters and Example Electrode Placement

Shown in Supplementary Fig. 2 is a picture of the parameters set on the DS7A neurostimulator during the experiment. An example of the electrode placement may be seen in Supplementary Fig. 3.

### Description of the Simulation Pipeline

Each run of the simulation began by specifying a total runtime (in seconds). At *t*_0_, to introduce noise, the initial membrane potential of all neuronal units was randomized between the reversal potential for the leak-current (*E*_*L*_, −80mV) and the threshold for action potential (*V*_*th*_, −50mV). The bladder was assumed to be empty (*V*_*B*_ = 0) with an initial inflow rate determined via a stochastic process. At each time step (*t*), the bladder state was updated and the firing rates of the primary pressure sensitive afferents (*F*(*P*_*B*_(*t*))) calculated. This firing rate was then used to update the status of the micturition control circuit and its efferent projections, which would be used to calculate the bladder state for (*t* + 1). The target values for the time step were then saved and the simulation increased to (*t* + 1). Upon reaching the final time step of the specified duration (*t*_*end*_) any cached data were collated and exported, and the simulation stopped.

### Synaptic Model

Synapses were modeled in terms of conductance using established formulae[42], modified to include opioidergic transmission. The magnitude of the synaptic current (*I*_*syn*_) was defined as:

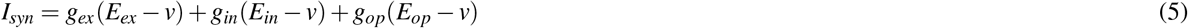

where *g*_*ex/in*_ represents the conductance of classical excitatory/inhibitory currents, and *E*_*ex/in*_ their reversal potentials; and *g*_*op*_/*E*_*op*_ representing the action of opioid currents. Modeled synaptic conductance decayed exponentially according to

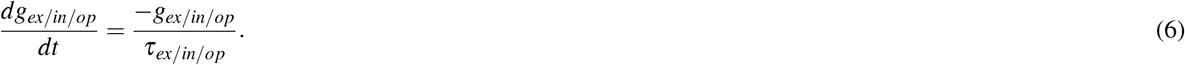

Synaptic strength was weighted, so that upon presynaptic spiking, the postsynaptic conductance of either excitatory, inhibitory, or opioidergic currents was incremented according to:

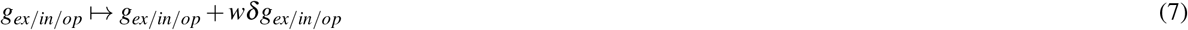

where *w* represents the weighting factor, and *δg*_*ex/in/op*_ the baseline increase in conductance in response to a presynaptic spike. The weights of all synaptic connections were determined by parameter fitting and constrained within a specific range to allow controlled plasticity within the network.

### Computational Model Optimization

Bayesian optimization was used to fit the parameters of the model [43]. To do so, the input to the micturition control circuit had to be carefully controlled. To this end, for the optimization stage, rather than implement a noisy biophysical model, the bladder was represented as a mathematical vector that contained experimentally derived data on internal bladder pressure (*P*_*B*_(*t*)) and volume (*V*_*B*_(*t*)). The sample rate of the vector was 50Hz, matching the model. The bladder data used to do so was obtained from previously published work[24] (see Animal Model Surgery and Preparation).

During optimization, the activity of second-order sensory afferents was extracted, downsampled to 10 Hz, and smoothed (1st-order Butterworth Filter, 1 Hz cutoff frequency). These data were compared to the ground-truth neural data associated with the input bladder data (see Animal Model Surgery and Preparation), producing a normalized root mean square error (NRMSE) value representing the difference between the simulated activity and expected activity. The algorithm adapted the weights and parameters of the model such that the NRMSE was minimized (acquisition function - Gaussian hedging, acquisition optimizer - LMBFGS, N_calls_ = 200). After running the algorithm, specific weights were manually adjusted to further optimize the efferent activity of the model, ensuring that it was accurate to typical bladder behavior.

Bayesian optimization of model weights and parameters considerably improved the accuracy of simulated afferent activity, bringing it closer to ground truth neural data (*NRMSE*_*baseline*_=3.367; *NRMSE*_*optimised*_=1.454). While subsequent manual tweaking did improve the efferent behavior of the model it did lead to a slight increase in NRMSE suggesting it did so at the cost of afferent accuracy (*NRMSE*_*fitted*_ = 1.599). Despite the minor decrease in global function fit, this methodology nevertheless improved the efferent behavior of the model and allowed the simulation to better capture the peaking pattern of neuronal firing rates in the afferent arm of the circuit (see Supplementary Fig. 1) providing an accurate foundation for analyzing the effects of neuromodulation. Notably, the optimization algorithm reached its minimum NRMSE plateau in a reasonable number of calls suggesting the global function minimum was true (see Supplementary Fig. 7).

Although the optimized model did not capture every peak in the firing rate or the physiological noise present in the data, the current level of accuracy is more than sufficient to serve as a foundation for exploring TNS, as noise omission would likely only affect the model’s ability to capture very specific network behaviors irrelevant to the present study.

### Model Parameters

Where possible these were matched to the parameters described in the original papers describing the neuronal, synaptic, and lower-urinary tract models[23, 27, 42]. Key parameters that were fit included the current mediating tonic activity (*I*_*ap*_), the magnitude of excitatory and inhibitory conductance (*g*_*ex/in/op*_), and the upper magnitude of spike-frequency adaption 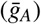. See Supplementary Table 1 for the parameters used in the simulation.

### Figure Design and Creation

Figures 1 and 2 were designed and created in BioRender. [44]

---

**Supplementary Figure 1.**
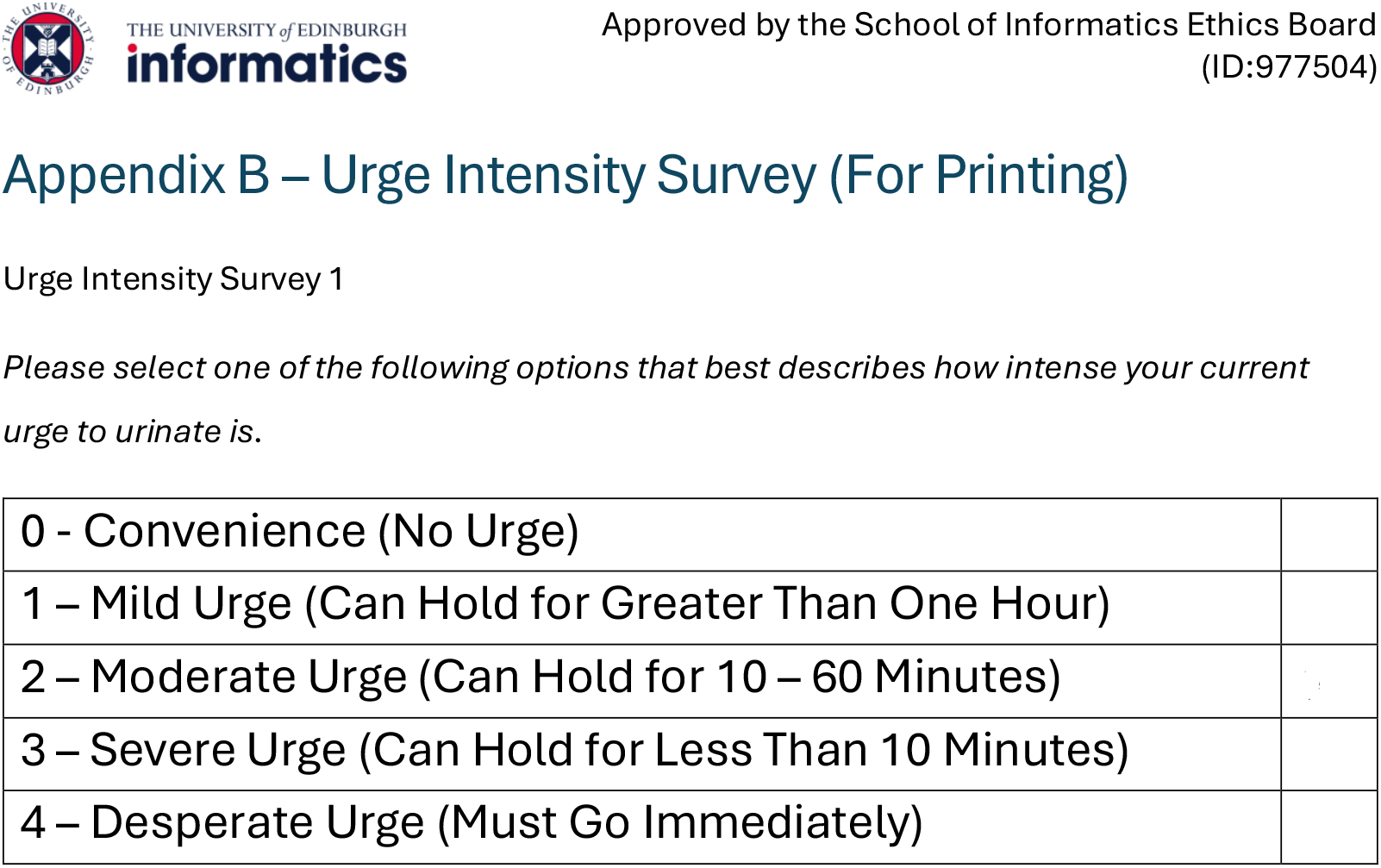
Example urge-intensity survey. Shown is an example of the survey which was given to participants in printed form.

**Supplementary Figure 2.**
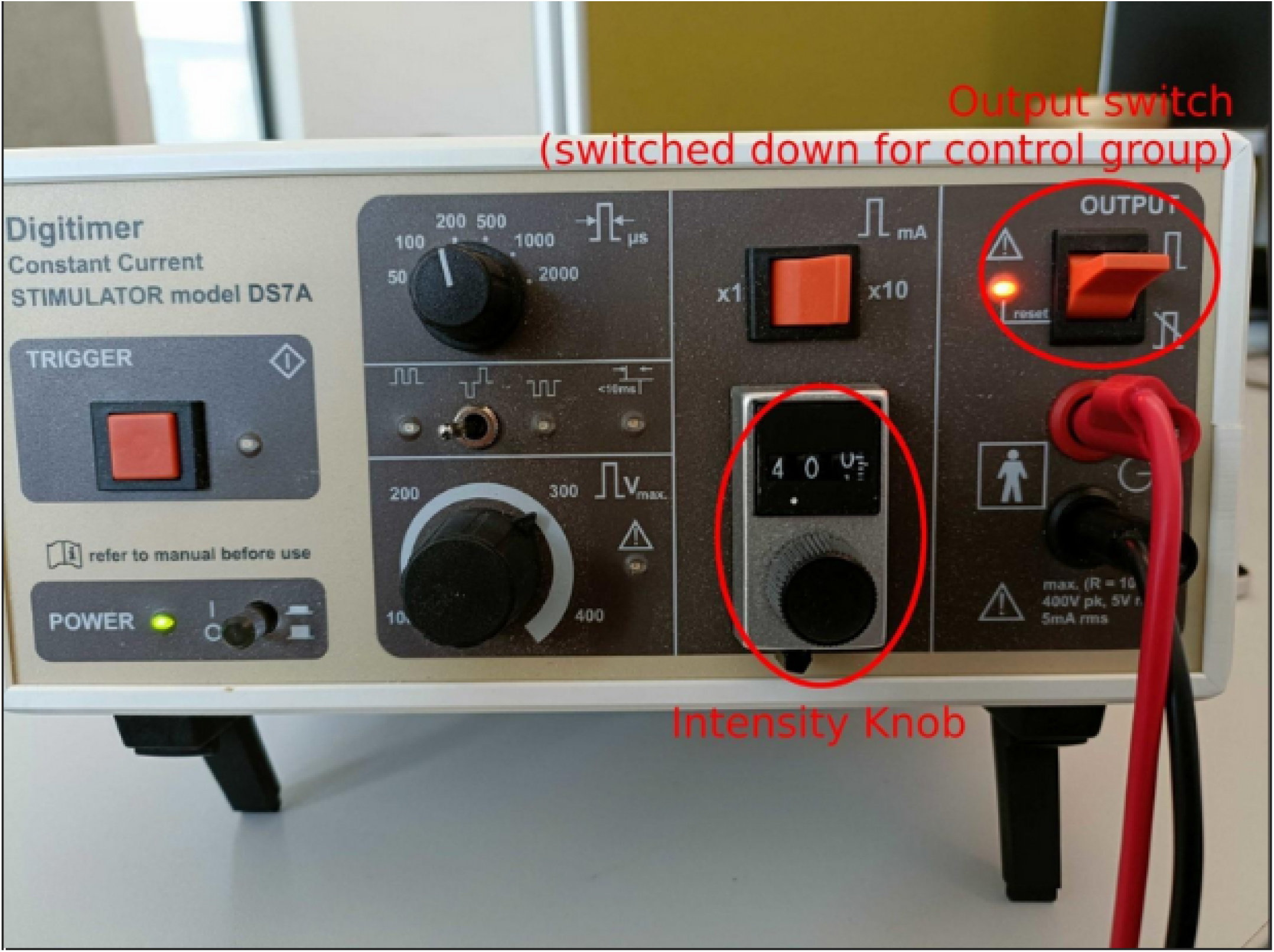
Parameters Used During Neurostimulation. Shown is the configuration of the DS7A neurostimulator as part of the experimental protcol. Stimulation was monophasic, with a 200*µs* pulse width, max-voltage of 300V, with a stimulation frequency that was set using a timing module built in-house that could output a trigger signal at 1 or 20 Hz.

**Supplementary Figure 3.**
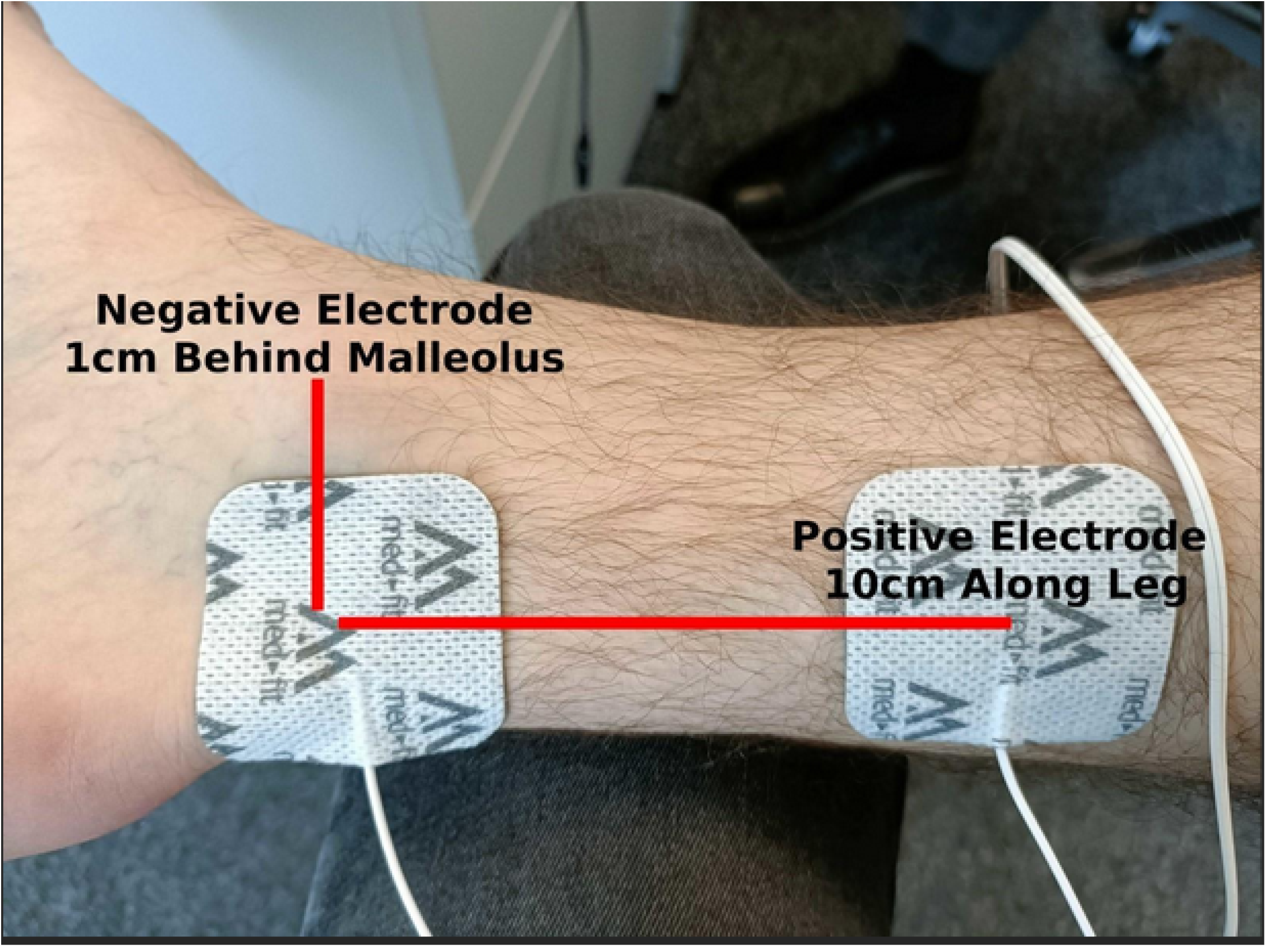
Standard Placement of TTNS Electrodes Used During Pilot Study.

**Supplementary Figure 4.**
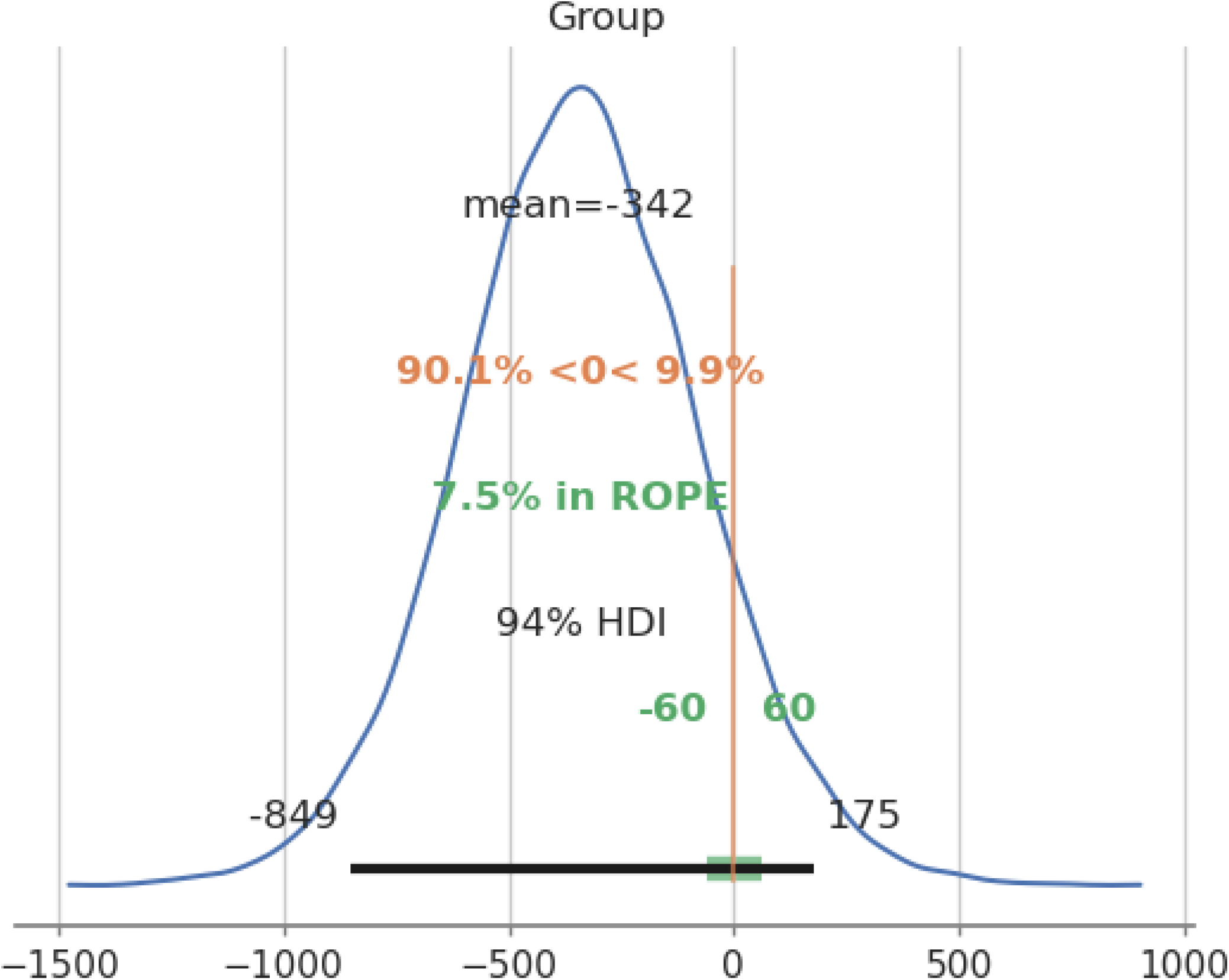
Pilot Study Time To First Urge ROPE Analysis. *N*_*draws*_ = 12, 000. Shown is the difference in time to first urge between groups A and B, in seconds.

**Supplementary Figure 5.**
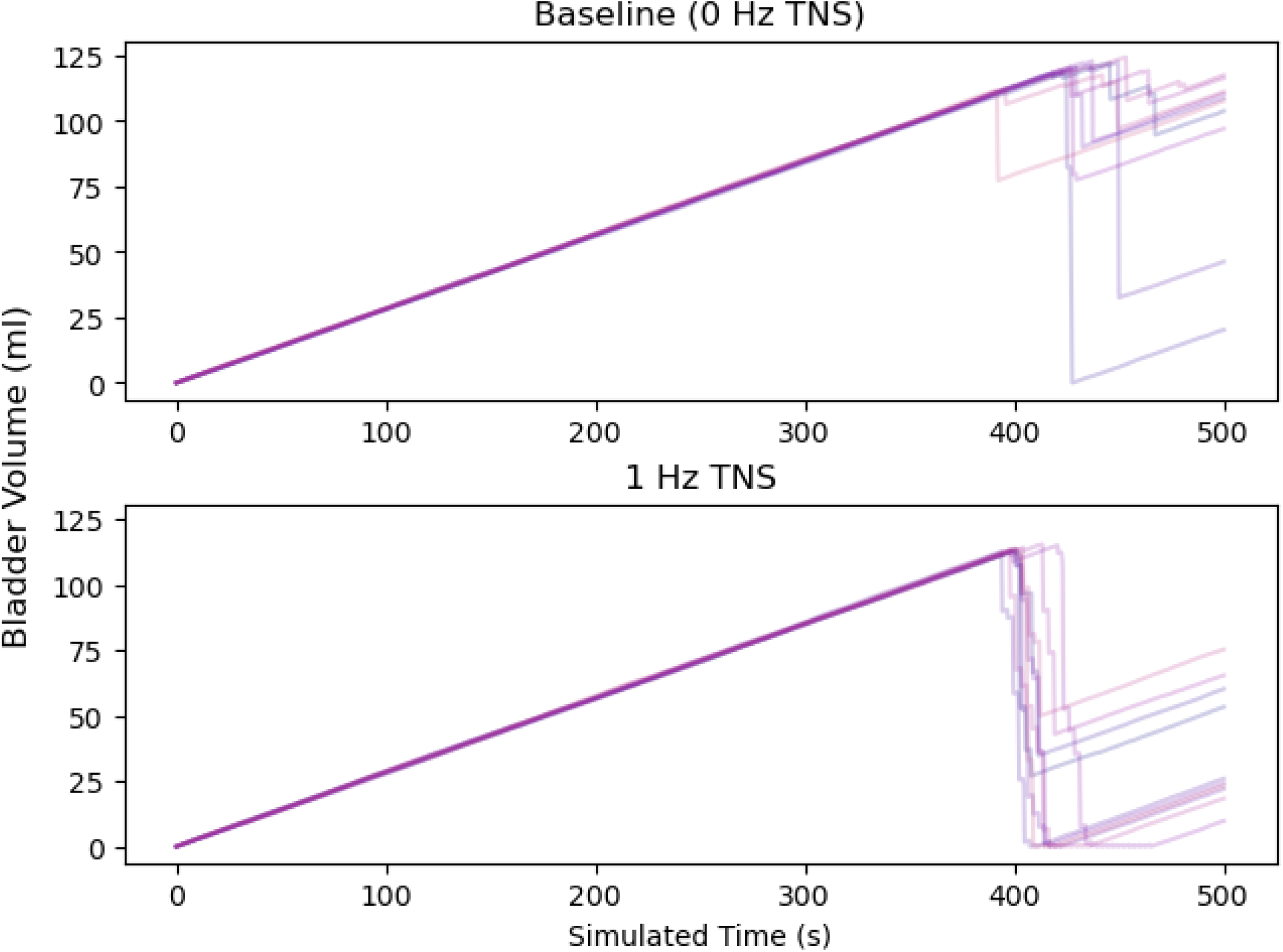
Simulated Bladder Volume Under Baseline and Low-Frequency Stimulation. Shown is the raw bladder volume from 10 independent bladder simulations (500s each) under baseline (0 Hz, top) and 1 Hz (bottom) tibial stimulation. All tibial nerve projections remained in-tact in both conditions.

**Supplementary Figure 6.**
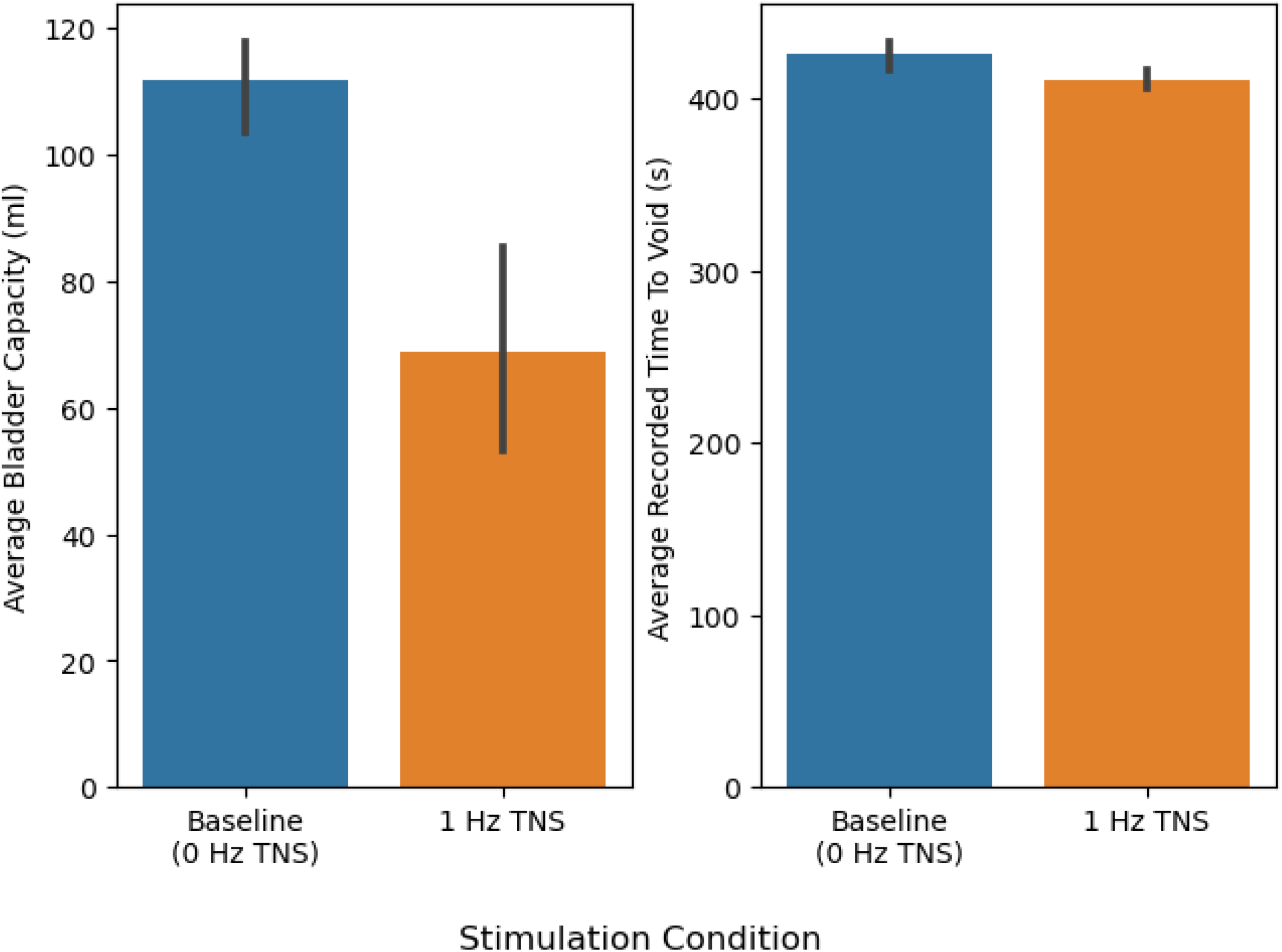
Impact of TNS on Bladder Capacity and Void Onset. Shown is the mean bladder capacity at the time of voiding (left) and the mean simulated time elapsed before voiding commenced (right) from 10 independent bladder simulations (500s each) under baseline (0 Hz) and 1 Hz tibial stimulation. All tibial nerve projections remained in-tact in both conditions. Error bars = 95% CI.

**Supplementary Figure 7.**
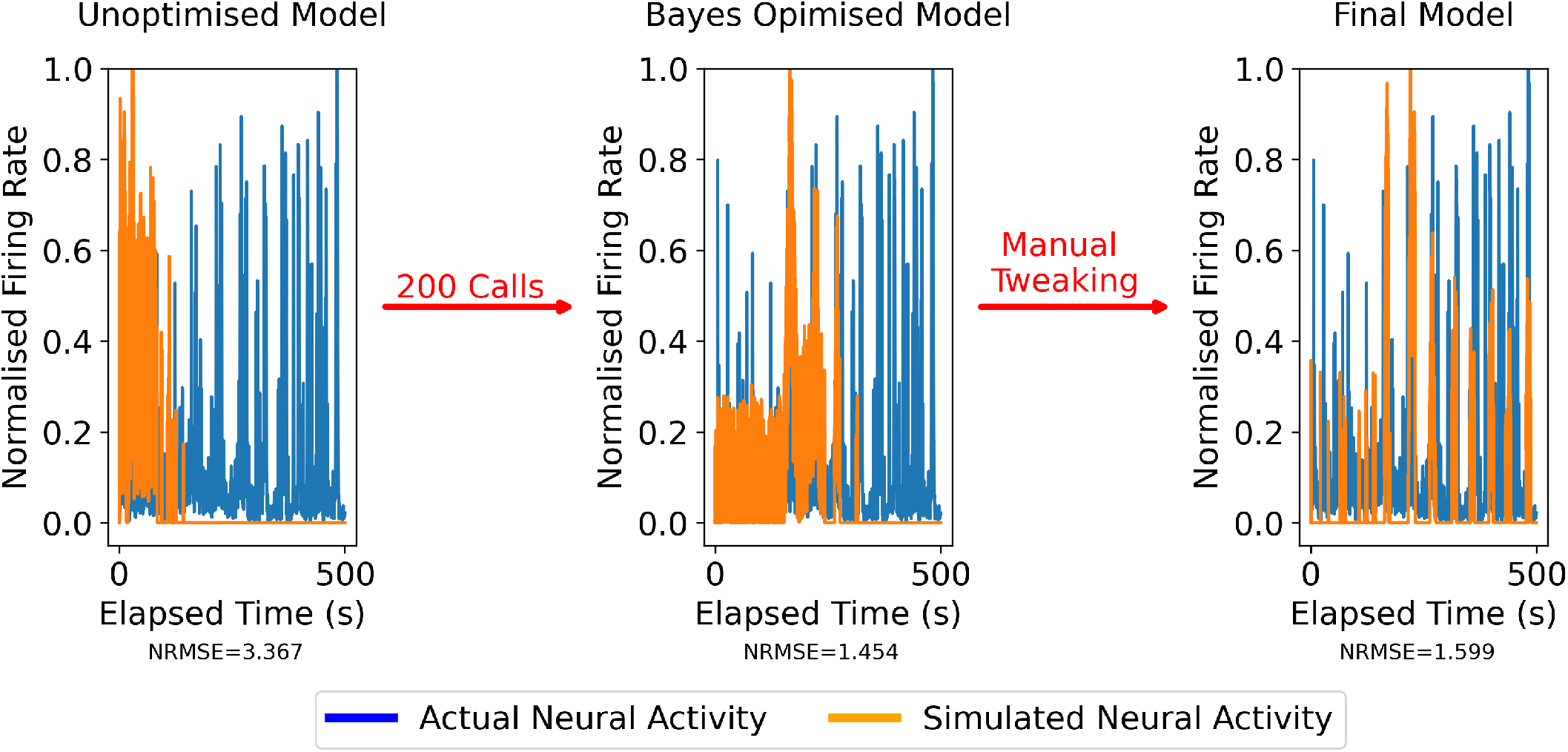
Model Performance During Fitting Process. Shown is the overlap between ground truth (blue) and simulated (orange) afferent neural activity for a subset of bladder pressure data. Model performance is shown in a random unoptimized state (left), after 200 rounds of Bayesian optimization (center), and after final manual optimization of model weights (right). The magnitude of the overlap between simulated and ground truth data at each stage shown as Normalized Root Mean Square Error (NRMSE). Smoothed firing rate (1st-order Butterworth filter, 1 Hz cutoff frequency) was normalized against the maximum recorded value in each dataset.

**Supplementary Figure 8.**
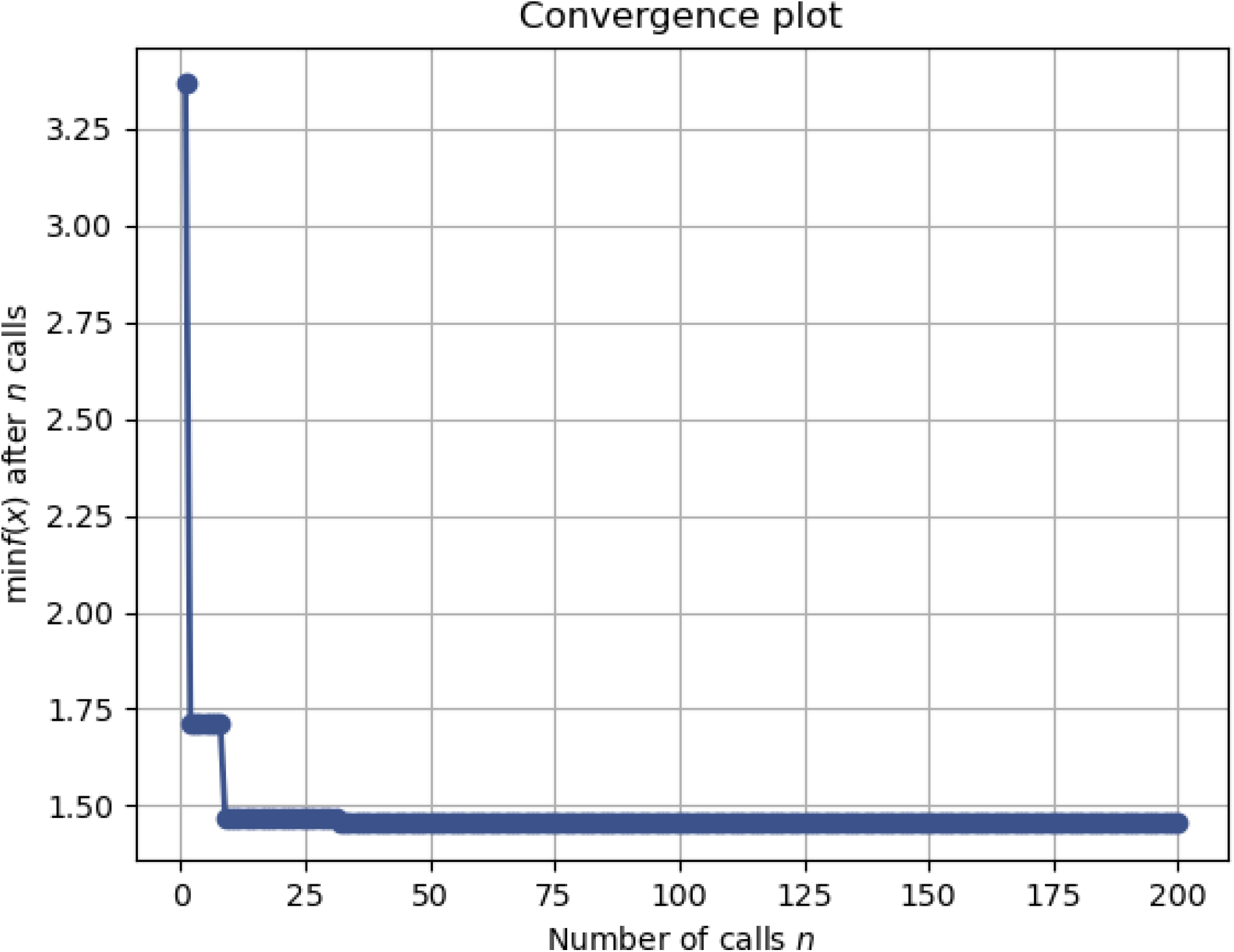
Bayesian Optimization Convergence Plot. Shown is a convergence plot detailing the Normalized Root Mean Square Error (NRMSE) of the ground truth vs. simulated afferent activity data at each phase of the optimization process.

**Supplementary Table 1.**
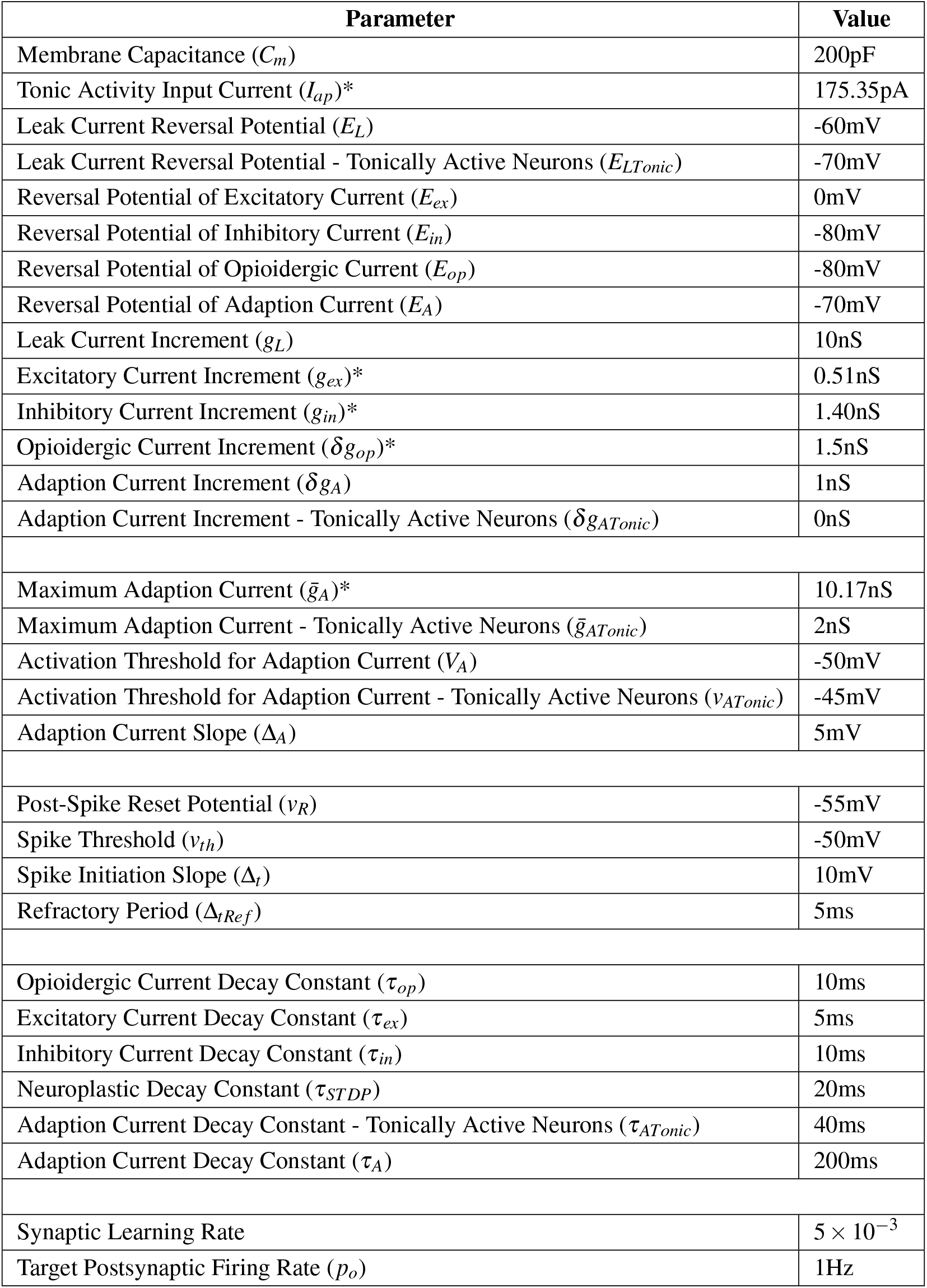
Final Neuronal Model Parameters. Where possible parameters were matched to the original specifications of the neuronal/synaptic model. Parameters that were altered by the model fitting process are marked as *.

## References

[1] Clare J. Fowler, Derek Griffiths, and William C. De Groat. “The neural control of micturition”. In: Nature reviews. Neuroscience 9.6 (June 2008), p. 453. ISSN: 1471003X. DOI: 10.1038/NRN2401.

[2] William C. de Groat, Derek Griffiths, and Naoki Yoshimura. “Neural Control of the Lower Urinary Tract”. In: Comprehensive Physiology 5.1 (Jan. 2015), pp. 327–396. ISSN: 20404603. DOI: 10.1002/CPHY.C130056.

[3] Daniel Jaskowak et al. “Mathematical modeling of the lower urinary tract: A review”. In: Neurourology and Urodynamics 41.6 (Aug. 2022), pp. 1305–1315. ISSN: 1520-6777. DOI: 10.1002/NAU.24995.

[4] William C. de Groat and Changfeng Tai. “Impact of Bioelectronic Medicine on the Neural Regulation of Pelvic Visceral Function”. In: Bioelectronic Medicine 2.1 (June 2015), pp. 25–36. ISSN: 23328886. DOI: 10.15424/BIOELECTRONMED.2015.00003.

[5] Tomonori Yamanishi et al. “Neuromodulation for the Treatment of Lower Urinary Tract Symptoms”. In: LUTS: Lower Urinary Tract Symptoms 7.3 (Sept. 2015), pp. 121–132. ISSN: 1757-5672. DOI: 10.1111/LUTS.12087.

[6] Mahipal Choudhary, Ron van Mastrigt, and Els van Asselt. “Inhibitory effects of tibial nerve stimulation on bladder neurophysiology in rats”. In: SpringerPlus 5.1 (Dec. 2016), pp. 1–8. ISSN: 21931801. DOI: 10.1186/S40064-016-1687-6.

[7] Thomas W. Fuller et al. “Sex difference in the contribution of GABAB receptors to tibial neuromodulation of bladder overactivity in cats”. In: American Journal of Physiology - Regulatory Integrative and Comparative Physiology 312.3 (Mar. 2017), R292–R300. ISSN: 15221490. DOI: 10.1152/AJPREGU.00401.2016.

[8] Todd Yecies et al. “Spinal interneuronal mechanisms underlying pudendal and tibial neuromodulation of bladder function in cats”. In: Experimental Neurology 308 (Oct. 2018), pp. 100–110. ISSN: 0014-4886. DOI: 10.1016/J.EXPNEUROL.2018.06.015.

[9] Zhaocun Zhang et al. “Role of µ, κ, and δ Opioid Receptors in Tibial Inhibition of Bladder Overactivity in Cats”. In: Journal of Pharmacology and Experimental Therapeutics 355.2 (Nov. 2015), pp. 228–234. ISSN: 0022-3565. DOI: 10.1124/JPET.115.226845.

[10] Yosuke Matsuta et al. “Contribution of opioid and metabotropic glutamate receptor mechanisms to inhibition of bladder overactivity by tibial nerve stimulation”. In: American Journal of Physiology - Regulatory Integrative and Comparative Physiology 305.2 (July 2013), pp. 126–133. ISSN: 03636119. DOI: 10.1152/AJPREGU.00572.2012.

[11] Matthew C. Ferroni et al. “Role of the brain stem in tibial inhibition of the micturition reflex in cats”. In: American Journal of Physiology - Renal Physiology 309.3 (Aug. 2015), F242–F250. ISSN: 15221466. DOI: 10.1152/AJPRENAL.00135.2015.

[12] Changfeng Tai et al. “Differential role of opioid receptors in tibial nerve inhibition of nociceptive and nonnociceptive bladder reflexes in cats”. In: American Journal of Physiology - Renal Physiology 302.9 (2012), pp. 1090–1097. ISSN: 15221466. DOI: 10.1152/AJPRENAL.00609.2011.

[13] Xunhua Li, Xing Li, and Limin Liao. “Mechanism of Action of Tibial Nerve Stimulation in the Treatment of Lower Urinary Tract Dysfunction”. In: Neuromodulation: Technology at the Neural Interface 27.2 (Feb. 2024), pp. 256–266. ISSN: 1094-7159. DOI: 10.1016/J.NEUROM.2023.03.017.

[14] Joanne Booth et al. “The effectiveness of transcutaneous tibial nerve stimulation (TTNS) for adults with overactive bladder syndrome: A systematic review”. In: Neurourology and Urodynamics 37.2 (Feb. 2018), pp. 528–541. ISSN: 1520-6777. DOI: 10.1002/NAU.23351.

[15] Alesha M. Sayner et al. “Transcutaneous Tibial Nerve Stimulation in the Management of Overactive Bladder: A Scoping Review”. In: Neuromodulation: Technology at the Neural Interface 25.8 (Dec. 2022), pp. 1086–1096. ISSN: 1094-7159. DOI: 10.1016/J.NEUROM.2022.04.034.

[16] Shun Li et al. “Prolonged nonobstructive urinary retention induced by tibial nerve stimulation in cats”. In: American Journal of Physiology - Regulatory Integrative and Comparative Physiology 318.2 (Feb. 2020), R428–R434. ISSN: 15221490. DOI: 10.1152/AJPREGU.00277.2019.

[17] Zainab Moazzam, Austin R. Duke, and Paul B. Yoo. “Inhibition and Excitation of Bladder Function by Tibial Nerve Stimulation Using a Wirelessly Powered Implant: An Acute Study in Anesthetized Cats”. In: The Journal of Urology 196.3 (Sept. 2016), pp. 926–933. ISSN: 15273792. DOI: 10.1016/J.JURO.2016.04.077.

[18] Katherine Theisen et al. “Frequency Dependent Tibial Neuromodulation of Bladder Underactivity and Overactivity in Cats”. In: Neuromodulation: Technology at the Neural Interface 21.7 (Oct. 2018), pp. 700–706. ISSN: 1094-7159. DOI: 10.1111/NER.12792.

[19] R. J. Maughan and J. Griffin. “Caffeine ingestion and fluid balance: a review”. In: Journal of Human Nutrition and Dietetics 16 (6 Dec. 2003), pp. 411–420. DOI: 10.1101/2024.11.21.624716.

[20] J H Burn, L H Truelove, and Isabel Burn. “THE ANTIDIURETIC ACTION OF NICOTINE AND OF SMOKING Professor of Pharmacology”. In: (1945).

[21] Jerry G. Blaivas et al. “The Urgency Perception Score: Validation and Test-Retest”. In: The Journal of Urology 177 (1 Jan. 2007), pp. 199–202. ISSN: 00225347. DOI: 10.1016/J.JURO.2006.08.091. URL: https://www.auajournals.org/doi/10.1016/j.juro.2006.08.091.

[22] W. C. De Groat and C. Wickens. “Organization of the neural switching circuitry underlying reflex micturition”. In: Acta Physiologica 207.1 (Jan. 2013), pp. 66–84. ISSN: 1748-1716. DOI: 10.1111/APHA.12014.

[23] Elliot Lister et al. “An Open-Source Neurodynamic Model of the Bladder”. In: BioRxiv (Nov. 2024). DOI: 10.1101/2024.11.21.624716.

[24] Milad Jabbari and Abbas Erfanian. “Estimation of Bladder Pressure and Volume from the Neural Activity of Lumbosacral Dorsal Horn Using a Long-Short-Term-Memory-based Deep Neural Network”. In: Scientific Reports 2019 9:1 9.1 (Dec. 2019), pp. 1–15. ISSN: 2045-2322. DOI: 10.1038/s41598-019-54144-8.

[25] Marcel Stimberg, Romain Brette, and Dan F.M. Goodman. “Brian 2, an intuitive and efficient neural simulator”. In: eLife 8 (Aug. 2019). ISSN: 2050084X. DOI: 10.7554/ELIFE.47314.

[26] Meredith J. McGee and Warren M. Grill. “Modeling the spinal pudendo-vesical reflex for bladder control by pudendal afferent stimulation”. In: Journal of Computational Neuroscience 40.3 (June 2016), pp. 283–296. ISSN: 15736873. DOI: 10.1007/s10827-016-0597-5.

[27] Tomasz Górski, Damien Depannemaecker, and Alain Destexhe. “Conductance-Based Adaptive Exponential Integrate-and-Fire Model”. In: Neural Computation 33.1 (Jan. 2021), pp. 41–66. ISSN: 0899-7667. DOI: 10.1162/NECO_A_01342.

[28] Walter F. Boron and Emile L. Boulpaep. Medical physiology. eng. Third edition. Philadelphia, PA: Elsevier, 2017. ISBN: 9781455733286.

[29] Jan Krhut et al. “Differences between brain responses to peroneal electrical transcutaneous neuromodulation and transcutaneous tibial nerve stimulation, two treatments for overactive bladder”. In: Neurourology and Urodynamics 42.6 (2023), pp. 1352–1361. DOI: 10.1002/nau.25197. eprint: https://onlinelibrary.wiley.com/doi/pdf/10.1002/nau.25197. URL: https://onlinelibrary.wiley.com/doi/abs/10.1002/nau.25197.

[30] Xunhua Li et al. “Real-time changes in brain activity during tibial nerve stimulation for overactive bladder: Evidence from functional near-infrared spectroscopy hype scanning”. In: Frontiers in Neuroscience Volume 17 - 2023 (2023). ISSN: 1662-453X. DOI: 10.3389/fnins.2023.1115433. URL: https://www.frontiersin.org/journals/neuroscience/articles/10.3389/fnins.2023.1115433.

[31] E. S. Lukacz et al. “A healthy bladder: a consensus statement”. In: International Journal of Clinical Practice 65 (10 Oct. 2011), p. 1026. ISSN: 13685031. DOI: 10.1111/J.1742-1241.2011.02763.X. URL: https://pmc.ncbi.nlm.nih.gov/articles/PMC3206217/.

[32] Mohamed S. Elkelini, Amal Abuzgaya, and Magdy M. Hassouna. “Mechanisms of action of sacral neuromodulation”. In: International Urogynecology Journal 21 (Oct. 2010), pp. 439–446. ISSN: 09373462. DOI: 10.1007/S00192-010-1273-3/FIGURES/2.

[33] Rosa L. Coolen et al. “Transcutaneous Electrical Nerve Stimulation and Percutaneous Tibial Nerve Stimulation to Treat Idiopathic Nonobstructive Urinary Retention: A Systematic Review”. In: European Urology Focus 7.5 (Sept. 2021), pp. 1184–1194. ISSN: 2405-4569. DOI: 10.1016/J.EUF.2020.09.019.

[34] Gabriele Gaziev et al. “Percutaneous tibial nerve stimulation (PTNS) efficacy in the treatment of lower urinary tract dysfunctions: A systematic review”. In: BMC Urology 13.1 (Nov. 2013), pp. 1–11. ISSN: 14712490. DOI: 10.1186/1471-2490-13-61.

[35] Aida Agost-González et al. “Percutaneous versus transcutaneous electrical stimulation of the posterior tibial nerve in idiopathic overactive bladder syndrome with urinary incontinence in adults: A systematic review”. In: Healthcare (Switzerland) 9 (7 July 2021). ISSN: 22279032. DOI: 10.3390/healthcare9070879. URL: /pmc/articles/PMC8306496/%20/pmc/articles/PMC8306496/?report=abstract%20 https://www.ncbi.nlm.nih.gov/pmc/articles/PMC8306496/.

[36] Manon te Dorsthorst et al. “Real-life patient experiences of TTNS in the treatment of overactive bladder syndrome”. In: Therapeutic advances in urology 13 (2021). ISSN: 1756-2872. DOI: 10.1177/17562872211041470. URL: https://pubmed.ncbi.nlm.nih.gov/34484428/.

[37] Allison S. Letica-Kriegel et al. “Identifying the risk factors for catheter-associated urinary tract infections: a large cross-sectional study of six hospitals”. In: BMJ Open 9 (2 Feb. 2019), e022137. ISSN: 2044-6055. DOI: 10.1136/BMJOPEN-2018-022137. URL: https://bmjopen.bmj.com/content/9/2/e022137 %20 https://bmjopen.bmj.com/content/9/2/e022137.abstract.

[38] Naglaa Youssef et al. “The Quality of Life of Patients Living with a Urinary Catheter and Its Associated Factors: A Cross-Sectional Study in Egypt”. In: Healthcare 11 (16 Aug. 2023), p. 2266. ISSN: 22279032. DOI: 10.3390/HEALTHCARE11162266. URL: https://pmc.ncbi.nlm.nih.gov/articles/PMC10454127/.

[39] Veronika Birkhäuser et al. “TASCI—transcutaneous tibial nerve stimulation in patients with acute spinal cord injury to prevent neurogenic detrusor overactivity: protocol for a nationwide, randomised, sham-controlled, double-blind clinical trial”. In: BMJ Open 10 (8 Aug. 2020), e039164. ISSN: 2044-6055. DOI: 10.1136/BMJOPEN-2020-039164. URL: https://bmjopen.bmj.com/content/10/8/e039164 %20 https://bmjopen.bmj.com/content/10/8/e039164.abstract.

[40] Wei Ju et al. “Smart Wearable TENS Device for Home-Based Overactive Bladder Management”. In: IEEE Transactions on Biomedical Circuits and Systems 19 (5 2025), pp. 981–992. ISSN: 19409990. DOI: 10.1109/TBCAS.2025.3527343.

[41] Aidan-MT/TibialModulationSim: A python based simulation of the bladder control circuit and the effects of tibial neuromodulation. URL: https://github.com/Aidan-MT/TibialModulationSim.

[42] T. P. Vogels et al. “Inhibitory plasticity balances excitation and inhibition in sensory pathways and memory networks”. In: Science 334.6062 (Dec. 2011), pp. 1569–1573. ISSN: 10959203. DOI: 10.1126/SCIENCE.1211095.

[43] Mike Diessner et al. “Investigating Bayesian optimization for expensive-to-evaluate black box functions: Application in fluid dynamics”. In: Frontiers in Applied Mathematics and Statistics 8 (Dec. 2022), p. 1076296. ISSN: 22974687. DOI: 10.3389/FAMS.2022.1076296.

[44] Scientific Image and Illustration Software | BioRender. URL: https://www.biorender.com/.

